# Molecular mechanism of α-latrotoxin action

**DOI:** 10.1101/2024.03.06.583760

**Authors:** BU Klink, A Alavizargar, KK Subramaniam, M Chen, A Heuer, C Gatsogiannis

## Abstract

The potent neurotoxic venom of the black widow spider contains a cocktail of seven phylum-specific latrotoxins (LTXs), but only one, α-LTX, targets vertebrates. This 130 kDa toxin binds to receptors at presynaptic nerve terminals and triggers a massive release of neurotransmitters. It is widely accepted that LTXs tetramerize and insert into the presynaptic membrane, thereby forming Ca^2+^-conductive pores, but the underlying mechanism remains poorly understood. LTXs are homologous and consist of an N-terminal region with three distinct domains, along with a C-terminal domain containing up to 22 consecutive ankyrin repeats. Here we report the first high resolution structures of the vertebrate-specific α-LTX tetramer in its prepore and pore state. Our structures, in combination with AlphaFold2-based structural modeling and molecular dynamics simulations, reveal dramatic conformational changes in the N-terminal region of the complex. Four distinct helical bundles synchronously rearrange to progressively form a highly stable 15 nm cation-impermeable coiled-coil stalk. This stalk, in turn, positions an N-terminal pair of helices within the membrane, thereby enabling the assembly of a cation-permeable channel. Taken together, these data unveil a unique mechanism for membrane insertion and channel formation, characteristic of the LTX family, and provide the necessary framework for advancing novel therapeutics and biotechnological applications.

## Main

Latrotoxins (LTXs) are the main toxic components of the venom from black widow spiders (*Latrodectus*)^1^. The venom includes the vertebrate-specific α-latrotoxin (α-LTX)^2,3^, five insecticidal toxins known as α, β, γ, δ, and ε-latroinsectotoxins (LITs)^4–6^, as well as a toxin specific to crustaceans named α-latrocrustatoxin (α-LCT)^7^. Upon envenomation, LTXs impact the victim’s nervous system by triggering massive neurotransmitter release upon binding to receptors^8–12^. α-LTX has been widely used as a molecular tool to study the exocytosis of synaptic vesicles and its actions are considered precisely the opposite of those of botulinum and tetanus toxins, both of which inhibit instead of activating the same secretory apparatus^13^.

LTXs are large proteins, ranging from 110 to 140 kDa, and they share a common architecture consisting of an N-terminal region containing functionally important cysteines and a C-terminal domain with up to 22 ankyrin repeats^14,15^. The mature toxins are created from non-toxic precursors by posttranslational cleavage of a short N-terminal and a larger C-terminal domain by Furin-like proteases^16^. In solution, LTXs exist as monomers or dimers^17–19^. However, in the presence of calcium (Ca^2+^) or magnesium ions (Mg^2+^), they have been observed to spontaneously oligomerize and integrate into the membrane, forming tetrameric pores that allow the selective passage of cations^18^. The efficiency of this process is significantly increased in the presence of receptors^20^. Recently, we reported cryo-EM structures of both the α-LCT monomer and the δ-LIT dimer, shedding light on the overall domain organization of the LTX family^19^. However, the mechanism of LTX pore formation remained elusive, as we currently lack detailed structures of a LTX tetramers and pore events have only been visualized at low resolution^18^. Understanding the mechanism of LTX action holds significant medical relevance^21,22^ and the potential to lead to the development of biotechnological applications and biopesticides^23^.

Here, we present the first high-resolution cryo-EM structures of the α-LTX tetramer in both the prepore and pore state. Combined with AlphaFold2 structural predictions and molecular dynamics simulations, we elucidate the long-sought-after α-LTX pore and unveil a unique mechanism characterized by dramatic conformational changes during α-LTX transition into a cation-selective channel.

### Cryo-EM structures of α-LTX

We used cryo-EM to image α-LTX from the mediterranean black widow spider (*Latrodectus tredecimguttatus*) in the presence of divalent cations to induce assembly into tetramers^18^. 2D classification revealed approximately 15% monomers, 19% dimers, 11% trimers and 55% tetramers (Supplementary figure 1a). The tetramers exhibited a preferred orientation with their symmetry axis exclusively perpendicular to the air-water interface, thus providing only a single view of the protein. An initial 2D analysis revealed the presence of at least two distinct tetrameric states, both displaying a characteristic central channel in a wide and narrow conformation (Figure 1a; Supplementary figure 1a; state 1 with 51 % and state 2 with 4% of particles, respectively). To address the preferred orientation issue, we collected multiple datasets with different tilt angles of the microscope stage from 0 to 60° and solved the structures of both states at 3.1 and 3.7 Å average resolution, respectively (Figure 1a,b; Supplementary figures 1-4, Supplementary table 1, Supplementary videos 1-2).

**Figure 1:**
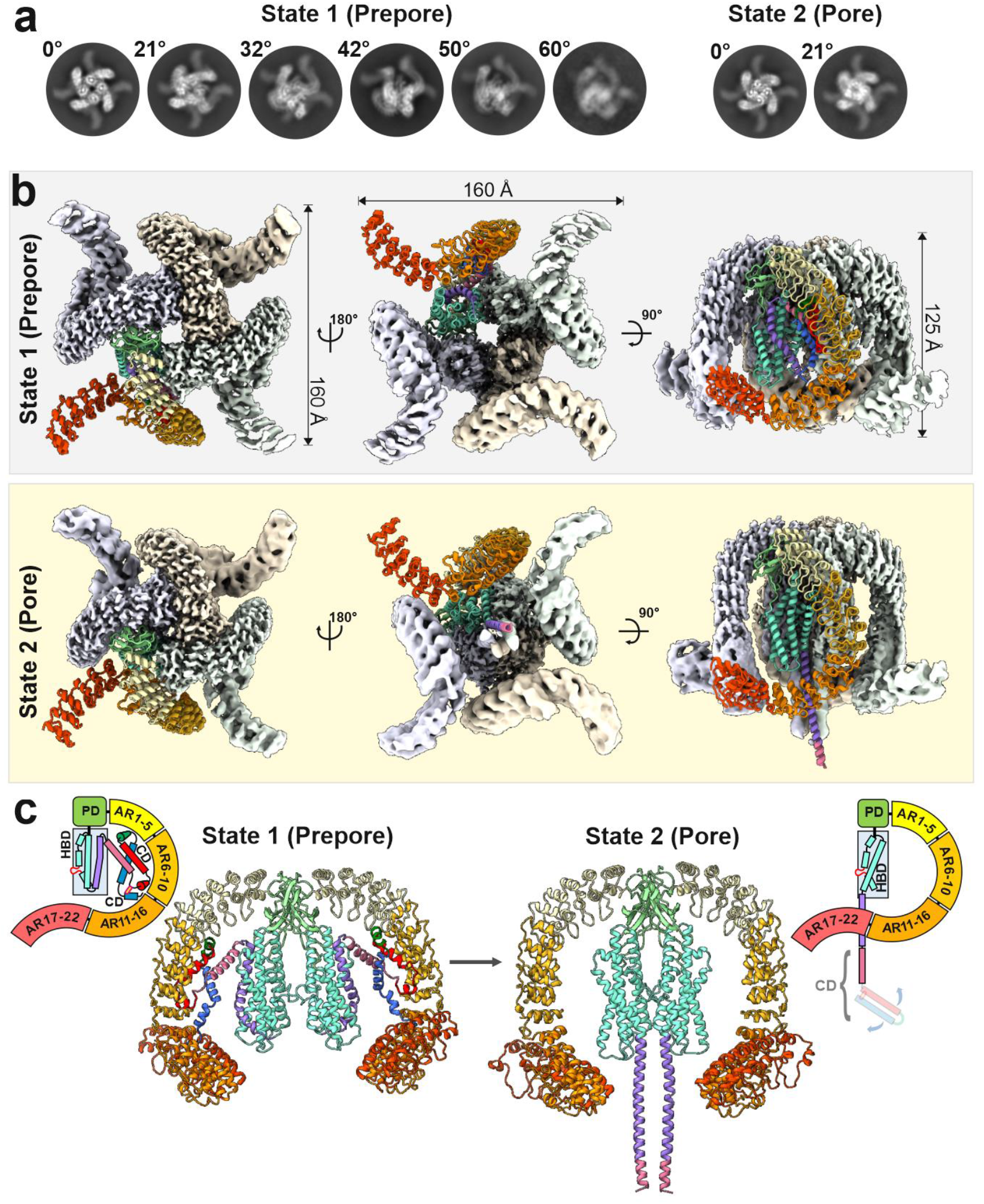
Cryo-EM structure of α-LTX in two distinct tetrameric states. **a** 2D class averages from datasets measured at different stage tilts. At low tilt angles, the presence of two distinct tetrameric states becomes apparent. Note the difference in diameter of the central channel. **b** Maps of the prepore (gray) and pore (yellow) state. For one monomer each, the derived molecular model is shown. **c** Molecular models of prepore and pore state of α-LTX. The side views depict two opposing subunits, colored according to their domain organization, as shown in the respective scheme. The first ∼100 residues of the CD (“tip of the needle”) are not resolved in the pore structure. CD: connector domain; HBD: helical bundle domain; ARD: ankyrin-like repeat domain; PD: plug domain.

The architecture of each subunit of state 1 has a characteristic G-shape and is similar to that of the soluble α-LCT monomer and the δ-LIT dimer^19^ (Figure 1b,c; Supplementary figure 5). Each monomer is composed of an N-terminal four-helix connector domain (CD; residues E21-D115), a central helical bundle domain (HBD; residues S116-K350), a short β-sheet plug domain (PD; residues Q351-D453) and a C-terminal tail of 22 ankyrin repeats (ARD; residues I454-G1195; Supplementary figure 5). Tetramerization does neither involve further large-scale conformational changes of individual subunits nor exposure of hydrophobic patches (compared to the soluble α-LCT monomer^19^), indicating that state 1 corresponds to the prepore state of α-LTX. The four HBDs form the core of the tetramer with a central ∼12-15 Å channel and the ARDs extend to the periphery, in a four-bladed windmill-like assembly (Figure 1b). The tetramer in the prepore state is assembled by stronger interactions between the PD of one monomer and ARs 1-6 of the clockwise neighboring monomer and is further stabilized by weaker interactions between the HBDs and PDs of neighboring monomers (Supplementary figure 6b,d). We observe significant movements of the HBDs and ARDs relative to each other, inducing conformation heterogeneities and symmetry breaks in the potential four-fold symmetry (Supplementary video 3). Interestingly, in the prepore state, the CD is sandwiched in a pocket between the three other domains of the monomer and completely avoids interactions with neighboring monomers (Figure 1c; Supplementary figure 5b).

In the second state, the HBDs are moved closer together, forming tight interactions that result in the narrow conformation of the central core (Supplementary figure 6c, f). The CD assumes a dramatically different conformation and is completely folded out of its pocket (Figure 1b,c). The four N-terminal CDs are remodeled in an extended coiled-coil needle, whereas the C-terminal domains preserve their overall arrangement, which results in a characteristic mushroom-like structure (Figure 1c), similar to other pore forming toxins^24–28^, indicating that state 2 represents the long-sought pore-state of α-LTX. The distal end of the tetrameric coiled-coil needle formed by the most N-terminal region of the protein (helices α1 to α3 in state 1) that might correspond to the transmembrane region of the pore is however flexible and not resolved in the cryo-EM structure, owing to the absence of a lipid environment or detergents.

### α-Latrotoxin inserts into membranes in an extended, mushroom-like conformation

It is well-established that α-LTX is able to integrate into biological membranes and this way forms cation-selective pores^20,29–32^. However, the underlying molecular mechanism and even the domains involved remained elusive. Despite significant efforts, we did not succeed to induce pore formation in presence of detergents, and reconstitution in lipid nanodiscs was too inefficient for subsequent cryo-EM studies of the pore. Once we added α-LTX to liposomes, we noticed by negative stain EM a small number of successfully reconstituted α-LTX particles (Figure 2a; Supplementary figure 7). This observation affirms α-LTX propensity for membrane insertion, but also aligns with previous studies suggesting that membrane insertion is highly inefficient in absence of receptors^20^. The particles show characteristic mushroom-like structures resembling the cryo-EM structure of state 2 with the (N-terminal) tip of their needle inserted into the membrane (Figure 2a,b). We therefore conclude that the mushroom-like cryo-EM structure of state 2 indeed represents the pore state of α-LTX.

**Figure 2:**
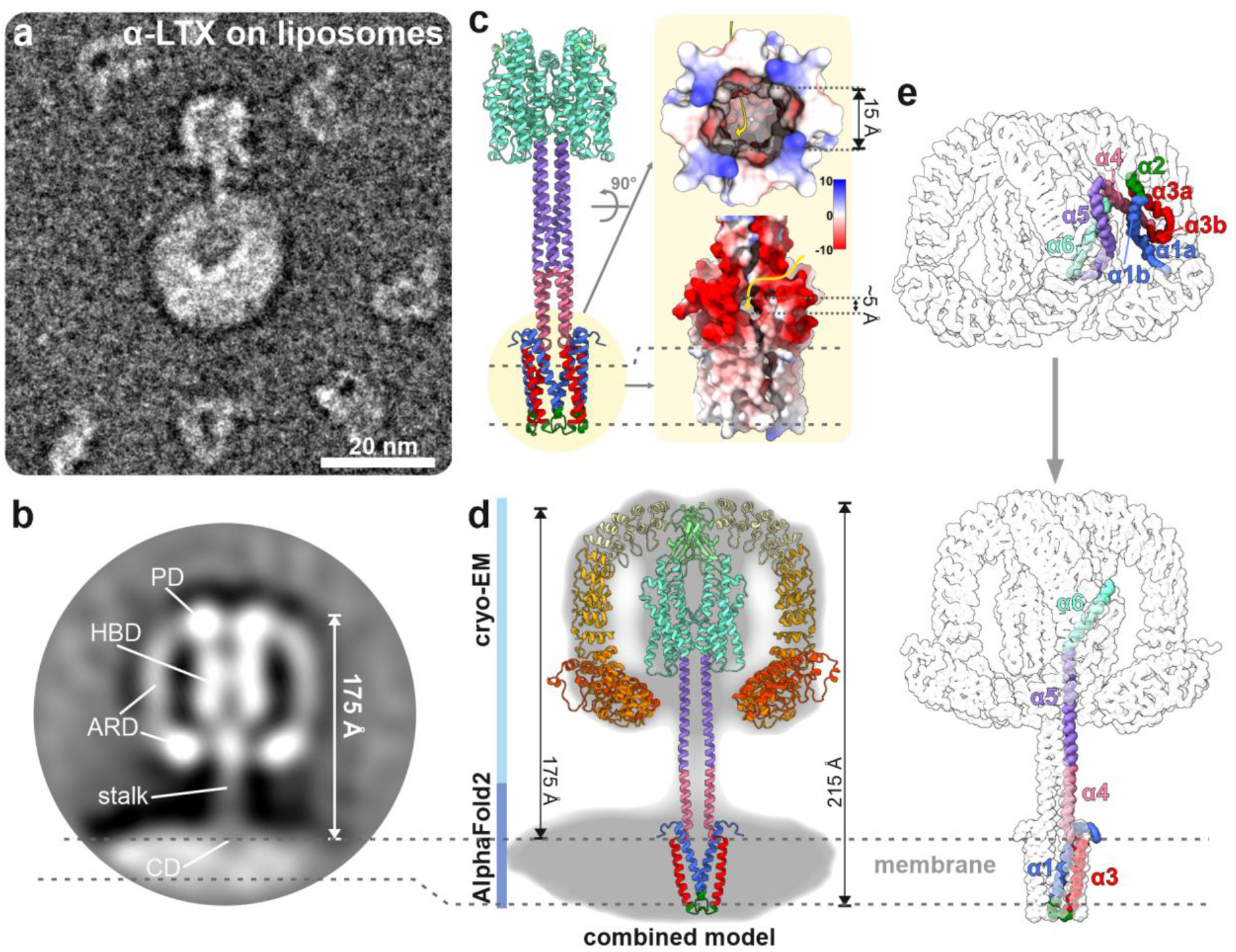
Molecular architecture of the α-LTX pore. **a**, α-LTX particle on a POPC-liposome in negative stain EM. **b** 2D average of 108 particles. The individual domains of the toxins are highlighted. **c** AlphaFold2 prediction of a tetrameric assembly of residues E21-E360 of α-LTX. The surface coulombic electrostatics [kcal/(mol·*e*)] indicates a membrane-spanning domain at its N-terminus. **d** Molecular model of the complete α-LTX pore obtained by combining the cryo-EM structure and the AlphaFold2 prediction overlaid in the negative stain average of α-LTX pore on liposomes. **e** Structural rearrangements of the CD leading to the formation of an extended coiled-coil and the transmembrane domain.

Using AlphaFold2, the tetrameric assembly of N-terminal residues E21-E360 (consisting of the central CD and HBD, in the absence of ARDs) was predicted, which complements the cryo-EM structure (residues F146-G1195) and also reveals the structure of the unresolved tip of the needle (Figure 2c; Supplementary Figure 8). In the overlapping region of the pore, the two structures agree very well (RSMD 0.9Å). While attempts to predict the structure of the full-length α-LTX tetramer using AlphaFold2 were not successful, we obtained a complete model of the α-LTX pore by combining the cryo-EM structure (residues F146-G1195) with the AlphaFold2 prediction of the missing region including the CD (residues E21-R145) (Figure 2d). These data allow a detailed mechanistic description of the pore formation events.

During prepore to pore transition, helix α5 separates from the HBD and undergoes a reorientation of almost 180 degrees (Figure 2e). This way, it forms an intriguing rigid tetrameric coiled-coil with the α5 helices of the other three monomers. In addition, helix α4 of the CD folds into a continuous helix with α5, this way elongating the coiled-coil and positioning N-terminal helices α1-α3 of the CD further away from the center of mass of the complex to form the transmembrane domain at the lower end of the coiled-coil (Figure 2e).

Helices α1-α3 thereby undergo dramatic conformational changes to form the transmembrane domain (TMD): In the prepore state, α1-α3 are isolated within each monomer, with a distance of ∼60-80 Å to each other, and are bent into a total of five shorter subhelices (α1a, α1b, α2, α3a, α3b; Figure 2e, Supplementary figure 5). In the pore state, they refold and arrange cylindrically into a disulfide-stabilized transmembrane bundle of 4×2 antiparallel helices formed by helices α1-α3 from all four monomers (Figure 2e). This way, a hollow barrel is assembled that is consistent with the expected features of a TMD of a pore forming toxin: While the assembly presents a highly hydrophobic patch spanning about two-thirds of its outside surface, it creates a negatively charged inner cavity with an average diameter of about 15 Å (Figure 2c). We performed MD simulations and the TMD remained stable in a POPC membrane supplemented with cholesterol and POPA (Supplementary figure 9; Supplementary video 4). The domain fully spans the lipid bilayer, while the 4×2 helix bundle itself is only penetrating with its hydrophobic patch. The outwards-facing third of the barrel is highly negatively charged, which acts as a stop signal, preventing further penetration of the stalk into the membrane (Figure 2c). Such charged “stop-patches” have also been observed for other pore-forming toxins^26,33^. In case of α-LTX, the partial integration of the 4×2 helical bundle leaves a ∼5 Å opening at the interface between the 4×2 helix bundle and the tetrameric coiled-coil (Figure 2c) which is framed by the negatively charged residues E28, E32, E38, D93 and D97 (Supplementary figure 10a).

### The cation-selective α-LTX channel

Even after identifying the transmembrane region within the CD, the translocation path of cations was at first glance not obvious. Interestingly, the HBDs, the tetrameric coiled-coil and the TMD form a ∼175 Å continuous channel (Figure 3a). Moreover, the HBDs provide a loop near the coiled-coil entrance, each displaying two aspartate residues (D209 and D210), which create a negatively charged pocket (Figure 3b). It was tempting to speculate that Ca^2+^ might enter the continuous channel through this site, in analogy to a “cation selectivity filter”, and subsequently pass through the coiled-coil towards the TMD. A similar mechanism has been recently proposed for Vip3 from *Bacillus thuringiensis*, a pore-forming toxin which also forms cation-selective pores^24^. Vip3 undergoes dramatic conformational changes upon proteolytic cleavage and similar to α-LTX, it forms a tetrameric coiled-coil stalk, even though the proteins have no further structural similarity (Supplementary figure 11). The authors identified a similar cation binding site close to the proximal entrance of the coiled-coil of Vip3a and therefore suggested that ions might get transported through its tetrameric stalk.

**Figure 3:**
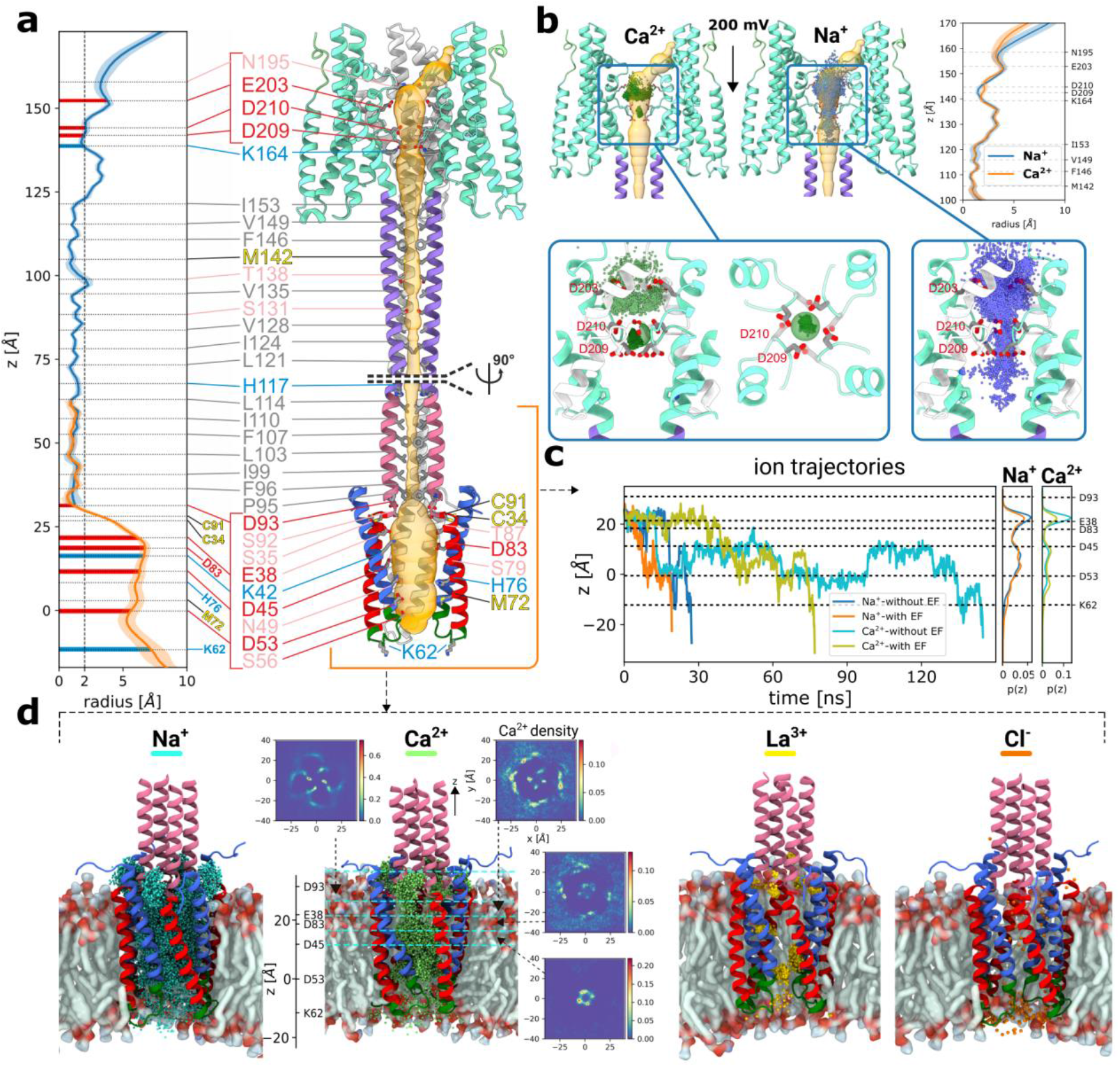
MD simulations of the pore structure of α-LTX. **a** Cartoon representation of the pore structure of α-LTX (residues E21-E360). The pore is represented as orange surface. Residues facing the pore volume are represented as sticks with labels colored according to their biochemical properties. The pore profiles from the two simulations (residues C91-Y260 for the upper part and the stalk; residues E21-S116 for the membrane protein part) are shown as blue and orange lines, respectively. **b** Cartoon representation of the Na^+^ and Ca^2+^ positions in the HBD, respectively, obtained from a superposition of many simulation snapshots with an applied electric field. Also, the resulting pore radius is shown. **c** Representative ion trajectories, corresponding to typical permeation events of the transmembrane domain by Na^+^ and Ca^2+^ ions, with and without an applied electric field. Additionally shown are ion density profiles obtained by averaging all successful permeation events. **d** Cartoon representations of simulations of the membrane part with different ions (Na^+^, Ca^2+^, La^3+^, Cl^-^) with a superposition analogous to **b**.The Ca^2+^ ionic densities in different slabs in the xy-plane are shown (5 Å and 10 Å widths along z, respectively).

In α-LTX, however, the radius of the cavity within the tetrameric coiled-coil is consistently less than 2 Å (Figure 3a), which is smaller than a Ca^2+^-oxygen binding distance and therefore too narrow for Ca^2+^ to pass through. Furthermore, MD simulations confirmed that this HBD pocket in α-LTX strongly binds Ca^2+^, but the affinity was so strong that even with an applied electric potential difference of 200 mV, a trapped cation was never released throughout the time frame of the simulation (Figure 3b). Instead, when a pulling force was applied to push the bound Ca^2+^ ion into the stalk, the cation quickly exited sideways (Supplementary figure 10b). Similarly, no Na^+^ was observed to enter the stalk (Figure 3b). Thus, it is highly unlikely that the stalk can transport ions, as further supported by its enormous stability, precluding any widening of its inner diameter (Supplementary figure 12).

Next, we addressed the putative cationic membrane transfer. We performed MD simulations of the TMD region and the distal end of the stalk in the presence of different ions. Na^+^ and Ca^2+^ ions were able to pass in large amounts; representative example trajectories are shown in Figure 3c. The higher charge of Ca^2+^ compared to Na^+^ has two different implications. First, due to the different Boltzmann factor, it results in higher selectivity when an electric field is applied. Second, Ca^2+^ has more localized positions than Na^+^ in the simulation, resulting in fewer and longer lasting membrane crossing events (Figure 3c,d; Supplementary tables 2-3; Supplementary figure 13; Supplementary video 5). These resulst are fully consistent with the reported properties of cation selective LTX pores *in vivo*^31,32,34*1*^. Cartoon representations of the simulation of the TMD (Figure 3d) clearly show the integrated positions of the cations during transport. By studying the height-dependent 2D-resolved Ca^2+^ density, it is even possible to identify the four entrance points and subsequent channels (Figure 3d). The entrance gate is localized at the 5 Å opening at the interface between the stalk and the TMD (Figure 2c; Supplementary figure 10a), which indeed fulfills important functions as a cation gate. La^3+^ ions are trapped at the entrance hole towards the cavity and block the pore (Figure 3d), in agreement with previous *in vivo* observations^*15*,*31*,*35*,*36*^. For Cl^-^, the strong negative charge of the entry region (Figure 2c, Supplementary figure 10a) is prohibitive. Taken together, our data reveal the gate of the α-LTX pore and fully explain its previously reported electrophysiological properties.

### Formation of tetramers prior membrane insertion

α-LTX purified in presence of Ca^2+^ consists of a mixture of all oligomeric prepore states from monomer to tetramer and a small population of the tetrameric pore assembly (Supplementary figure 1a). With the existence of trimeric prepore species, we can exclude the notion that two α-LTX dimers are required for the formation of a tetrameric prepore, as previously proposed^37^. A stepwise assembly of the tetrameric prepore can easily be rationalized by the involved subunit interactions, which are almost exclusively formed between the PD of one monomer with the AR domain of the clockwise neighboring monomer (Supplementary figure 6).

Tetramerization was blocked in the N4C-mutation of α-LTX by insertion of residues VPRG before the first AR^15^. This mutant abolishes its ability to form pores and is therefore extensively used in the field to selectively study the receptor-mediated actions of α-LTX^17,38–40^. The high-resolution structure of the tetrameric prepore shows that the introduced VPRG residues are placed N-terminal of the last ß-sheet (ß7) of the PD domain, disturbing the crucial intermolecular oligomerization site between PD and neighboring ARD (Supplementary figure 14). This explains why this mutant loses its ability to oligomerize and form ionophores^17,36^, while it retains the ability to bind and activate receptors^17,40^.

Consistent with previous functional studies^37^, our data support the notion that the formation of the tetrameric prepore assembly is a prerequisite for the formation of the pore state (Supplementary figure 6a). The dramatic prepore to pore rearrangement relies mostly on the successful formation of the rigid tetrameric coiled-coil by helices α4 and α5. The stability of the coiled-coil is quantified by studying the strong loss in free energy when slowly removing one monomer of the prepore structure as reflected by an increasing RMSD as compared to the tetrameric coiled-coil structure in the pore state (Supplementary figure 12). Moreover, the interactions within the tetrameric coiled-coil in the pore state are about an order of magnitude stronger than the PD/ARD interactions, which have the strongest contribution in the prepore state (∼30 vs ∼3.3 kcal/mol, respectively) (Supplementary table 4).

Importantly, only towards formation of the pore state, the HBDs move closer together to form several stabilizing contacts (Figure 1b; Supplementary figure 6c,f), including the prominent Ca^2+^ binding site directly above the coiled-coil stalk (Figure 3b). This site binds Ca^2+^ with such high affinity that it cannot easily be released (Figure 3b; Supplementary figure 10b), suggesting that Ca^2+^ binding supports the dramatic conformational rearrangement of the prepore tetramer into the pore state. Furthermore, also the observation that the cavity radius in the upper part of the HBD pocket is smaller for Ca^2+^ than for Na^+^ (Figure 3b) may suggest a Ca^2+^ stabilizing mechanism for the pore state.

### Role of disulfide bridges for the transition to the pore state and receptor binding

Previous biochemical studies suggest that disulfide bond reduction abolishes the activity of α-LTX and its binding to receptors^11,15^. α-Latrotoxin contains a total of 9 cysteine residues, two of which are in the N-terminal CD, one in the PD and six in the C-terminal AR15-19 (Figure 4a).

**Figure 4:**
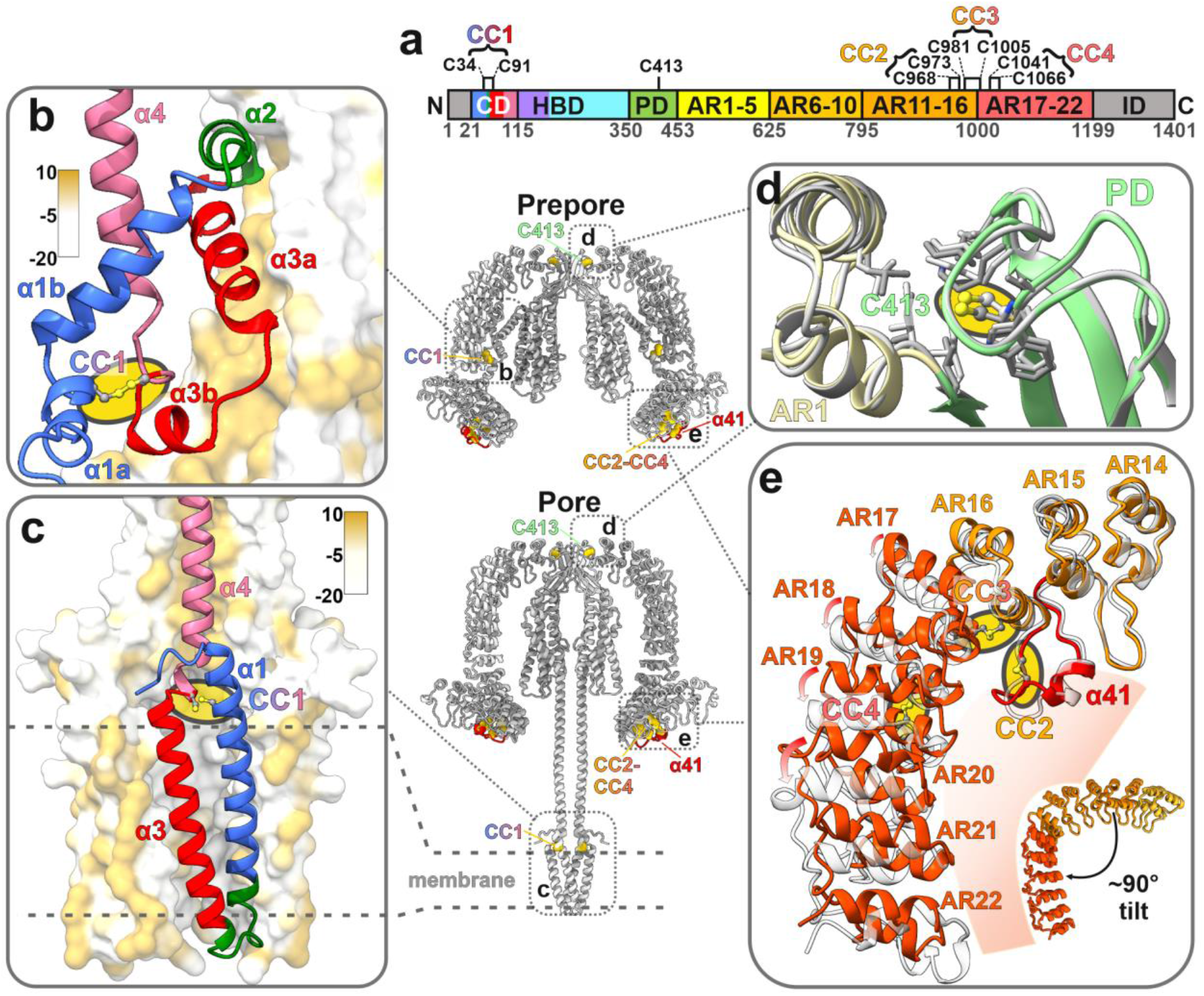
α-LTX stabilization by cysteines and disulfide bonds. **a**, Position of cysteines in α-LTX. Disulfide pairs are marked with braces. **b**, CD in the prepore state. The interacting AR domain is contoured by its surface hydrophobicity. **c**, CD in the pore state. The interacting CDs from different α-LTX monomers are contoured by surface hydrophobicity. **d**,**e** Cys413 and disulfide bonds in the AR domain. The prepore state is colored and the pore state in light grey.

The N-terminal conserved cysteines of the CD were mutated to serine (C34S, C91S and C413S) and were found to be essential for α-LTX action^15^. Our data confirm that the conserved C34 and C91 are involved in a stabilizing disulfide bridge (CC1) between helices α1 and α3 of the CD, both in the prepore and the pore states (CC1; Figure 4b,c). It should be emphasized that these helices form the membrane-spanning α-LTX channel (Figure 2e). Although the fold of helices α1-α3 is remarkably different between the two states (Figure 4b,c), CC1 is a crucial stabilizing connection for this assembly and thus for the prepore to pore transition, as CC1 stays intact during the refolding and massive reorientations of up to ∼190 Å from prepore to pore state. Alphafold2 predictions further confirm that the stabilization of the membrane-spanning domain by CC1 is conserved throughout the LTX family (Supplementary figure 15a).

In contrast, C413 does not form a disulfide bond but is a central part of a hydrophobic pocket connecting PD and ARD in α-LTX (Figure 4d). Loss of function upon mutation of C413 to a less hydrophobic serine is thus not due to lack of disulfide potential, but rather due to altered protein fold.

Furthermore, our structures reveal that the six C-terminal cysteines in AR15-AR19 of α-LTX form three disulfide bonds (CC2, CC3, CC4), which might be important for receptor binding and toxin specificity (Figure 4e; Supplementary figure 15), since they are located in the region that contains the interaction site for the receptor Latrophilin^41,42^. Removal of ARs 15-22 containing CC2, CC3 and CC4 results in a complete loss of α-LTX stimulating activity^43^.

CC2 is the first disulfide bond in the ARD (C968-C973) in α-LTX and is located within an extended loop between ARs 15 and 16, succeeding the short helix α41 located within this loop (Figure 4e). This extended loop between AR 15 and 16, in combination with an unusually short loop between AR16 and AR17, introduces a rotation in the alignment of ARs to each other by almost 90°. CC3 and CC4 in α-LTX are located directly after the extended loop between ARs 15 and 16, which qualifies them to stabilize this characteristic bending point in the middle of the ARD tail (Figure 4e).

Interestingly, the position of the characteristic loop and CC2 in the AR tail are conserved in the vertebrate-specific α-LTXs, but have a different arrangement in the crustacean-specific α-LCT and the insect-specific α-LIT. These do not contain such an extended loop between AR 15 and 16, but a loop inserted within AR18. This loop is also stabilised by a disulfide bond, which we call CC2* in α-LCT and α-LIT for better clarity. According to AlphaFold2 predictions, this distinct positioning of the disulfide-stabilised loop also results in a different bending of the AR tail (Supplementary Figure 15b). In addition, the insect-specific δ-LIT contains only 13 ARs, i.e. its tail is truncated and does not contain these disulfide bonds^19^, but instead a highly negatively charged domain that could represent an alternative receptor binding motif (Supplementary Figure 15b).

The ARDs containing the receptor binding sites are otherwise conserved across the members of the LTX family, known for being highly phylum-specific. It is conceivable that the consistent positioning of disulfide bonds CC3-4, coupled with variable positions of the loop containing CC2 or CC2*, plays a crucial role in receptor binding. The architecture of the loop, along with the curvature and length of the ARD tail, emerges as the most likely set of factors determining LTX target specifity.

## Discussion

Our study elucidates the process through which α-LTX integrates into presynaptic membranes and creates a selective cation-permeable channel that facilitates calcium influx, triggering uncontrolled neurotransmitter release. The results presented here explain numerous long-standing questions regarding the mechanism of LTX action accumulated during the last decades. Based on our findings and previous work in the field, a model for α-LTX action at the presynaptic membrane is presented in Figure 5.

**Figure 5.**
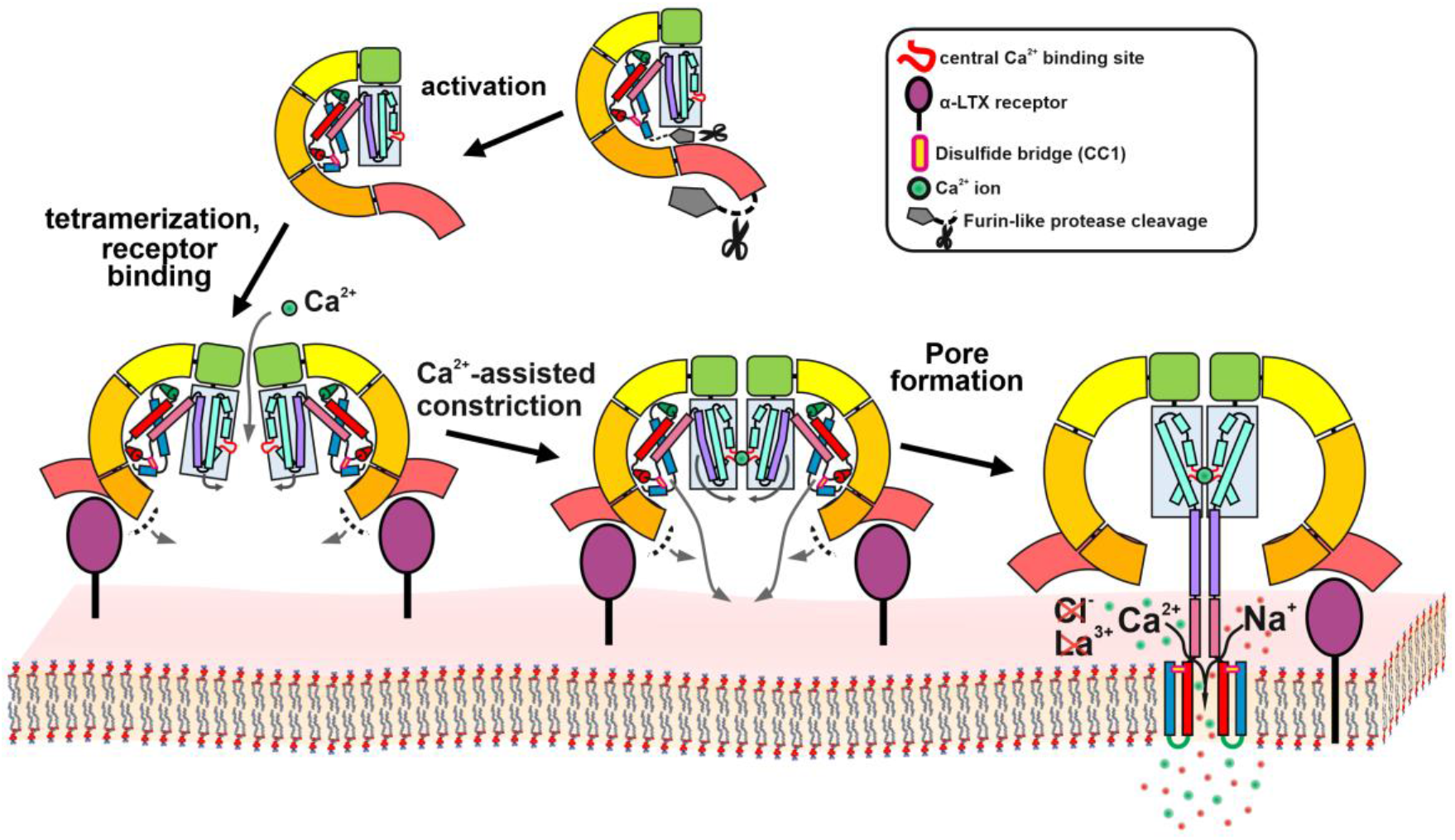
Schematic model of α-LTX activation, tetrameric prepore assembly and transition to a cation permeable channel.

An α-LTX precursor is cleaved by furin-like proteases at its N- and C-termini, producing an activated soluble G-shaped toxin monomer^16^. Four monomers assemble stepwise in a clockwise manner via interactions between the PD and ARD domains to form the tetrameric prepore state that is competent to initiate membrane-insertion. This process is most probably supported by binding of receptors to the ARD-tails, which also brings the complex in proper orientation and close proximity to the membrane. Noteworthy, the formation of pores is possible in pure lipid membranes^20,32^ (Figure 2a), but the presence of receptors significantly increases the occurrence of pores^20^.

The prepore displays a high degree of conformational flexibility in the curvature and/or relative orientation of different subunits and domains, resulting in “breathing motions” of the central channel (Supplementary Video 6). We believe that narrowing of the channel, as seen in subsets of our cryo-EM data, marks a first prerequisite step to initiate pore formation: only within a sufficiently narrow channel, the four loop regions containing aspartates D209 and D210 are close enough to form a central high-affinity cation binding pocket as seen in the pore state (Figure 3b; Supplementary Video 6). This process, which most probably progresses in a stepwise manner, also pulls the termini of helices α6 close enough together to initiate the formation of the tetrameric stalk. It is conceivable that the stabilization of the narrow conformation by Ca^2+^ ions in the D209/D210 binding pocket is the determining factor for the described Ca^2+^-dependence of α-LTX pore formation^35,37^. Moreover, in the prepore state, the helices of the N-terminal CD are tightly packed and protected within a groove formed between the HBD and the ARD-tail, with helices α1-α3 being fragmented, but stabilized by a conserved disulfide bond. Ca^2+^ dependent narrowing of the central HBD channel, combined with the tetrameric stalk formation starting at the hinge between helices α6 and α5, destabilizes the groove harboring the CD.

Upon opening of the groove, the assembly of the tetrameric stalk can now proceed by spontaneous rearrangements of the CDs elongating the prearranged N-termini of the tetrameric coiled-coil stalk, up to its final length of about 15 nm. This process might resemble the assembly of SNARE complexes which promote vesicle fusion by the formation of a highly stable tetrameric coiled-coil^44^. The four N-terminal α-helical pairs of the CD, each stabilized by the conserved disulfide bond that remains intact during this conversion, come close to each other at the low end of the stalk. Together the CDs insert into the membrane to assemble the transmembrane channel. The prepore-to-pore transition converts the flat windmill-like prepore complex into an extended mushroom-like pore conformation (Figure 5; Supplementary videos 7-8).

Notably, the HBD, the stalk and the transmembrane domain thereby form a continuous channel. The stalk is however not permeable for ions but rather functions as a molecular ruler: α-LTX receptors exhibit an elongated shape, with the extracellular domain protruding far from the membrane^45,46^. Upon receptor binding, the coiled-coil stalk possesses an appropriate length to span this distance and deliver the CD to the presynaptic membrane (Supplementary Video 7). In light of the substantial height of the pore state of approximately 175 Å protruding from the membrane, these dimensions remain in accord with the ∼20 nm width of the synaptic cleft^47^.

A side-entry gate, selective for monovalent and divalent cations, is located at the interface between the coiled-coil stalk and the transmembrane CD. Ca^2+^ ions enter this gate from the side of the stalk directly above the upper leaflet of the presynaptic membrane and pass through the transmembrane pore (Supplementary Video 7). This mechanism mimics Ca^2+^ influx via voltage-gated Ca^2+^ channels during an action potential and triggers massive synaptic vesicle exocytosis.

Our data allows understanding of the α-LTX mechanism of action at an unprecedented level of detail. α-LTX stands out as an exceptional pore-forming toxin (PFT), functioning differently from any other known toxin. The mechanistic understanding of α-LTX function provides a strong foundation towards the development of novel biomedical applications, including treatment of latrodectism and botulism, improved tools for studying neurotransmitter exocytosis, and the development of specific bio-insecticides.

## Data availibility

The cryo-EM maps of “prepore” and “pore” state have been deposited to the Electron Microscopy Data Bank (EMDB) under the accession codes [EMD-XXX](https://www.ebi.ac.uk/emdb/EMD-16372) and EMD-XXX. The cryo-EM dataset has been deposited to EMPIAR under accession codes [EMPIAR-XXX](https://dx.doi.org/10.6019/EMPIAR-11671). The coordinates of the corresponding models have been deposited to the Protein Data Bank (PDB) under accession codes [XXX](https://www.rcsb.org/structure/8C0V) and XXX. Other data are available from the corresponding author upon request.

## Acknowledgements

The cryo-EM data were collected at “Cryo-EM SoN”, the cryo-EM infrastructure of the University of Münster, funded by the Deutsche Forschungsgemeinschaft (DFG, German Research Foundation) – Projectnumber 496113311. We thank Dr. Neuhaus and Dr. Blanque (cryo-EM SoN, Center for Soft Nanoscience) for technical support. The cryo-EM dataset was processed at the Palma II HPC (DFG INST 211/667-1) of the University of Münster. The MD simulations have also been carried out on the Palma II cluster. We acknowledge technical support by the HPC team of the University of Münster. This work was supported by the DFG (SFB1348 to A.H. and C.G.).

## Author contributions

C.G. designed and managed the project. B.U.K. collected and processed cryo-EM data with contributions by M.C. at the initial stage of the project. B.U.K. analyzed cryo-EM data, performed AF2 predictions, built atomic models and interpreted data with contributions by C.G. K.K.S., M.C. performed liposome reconstitution experiments, negative stain EM screenings and processed negative stain EM data. A.A. performed MD simulations. A.A and A.H. analyzed MD data. B.U.K., C.G. drafted the manuscript. B.U.K., A.A., A.H., C.G wrote the manuscript. All authors discussed the results, commented and approved the manuscript.

## Ethics declarations

The authors declare no competing interests.

