## Supplement for "Molecular mechanism of α-latrotoxin action"

# of

**Contents:**

1. Supplementary Figures

2. Supplementary Tables

3. Supplementary Videos

4. Supplementary References

### 1. Supplementary Figures

#### Supplementary figure 1

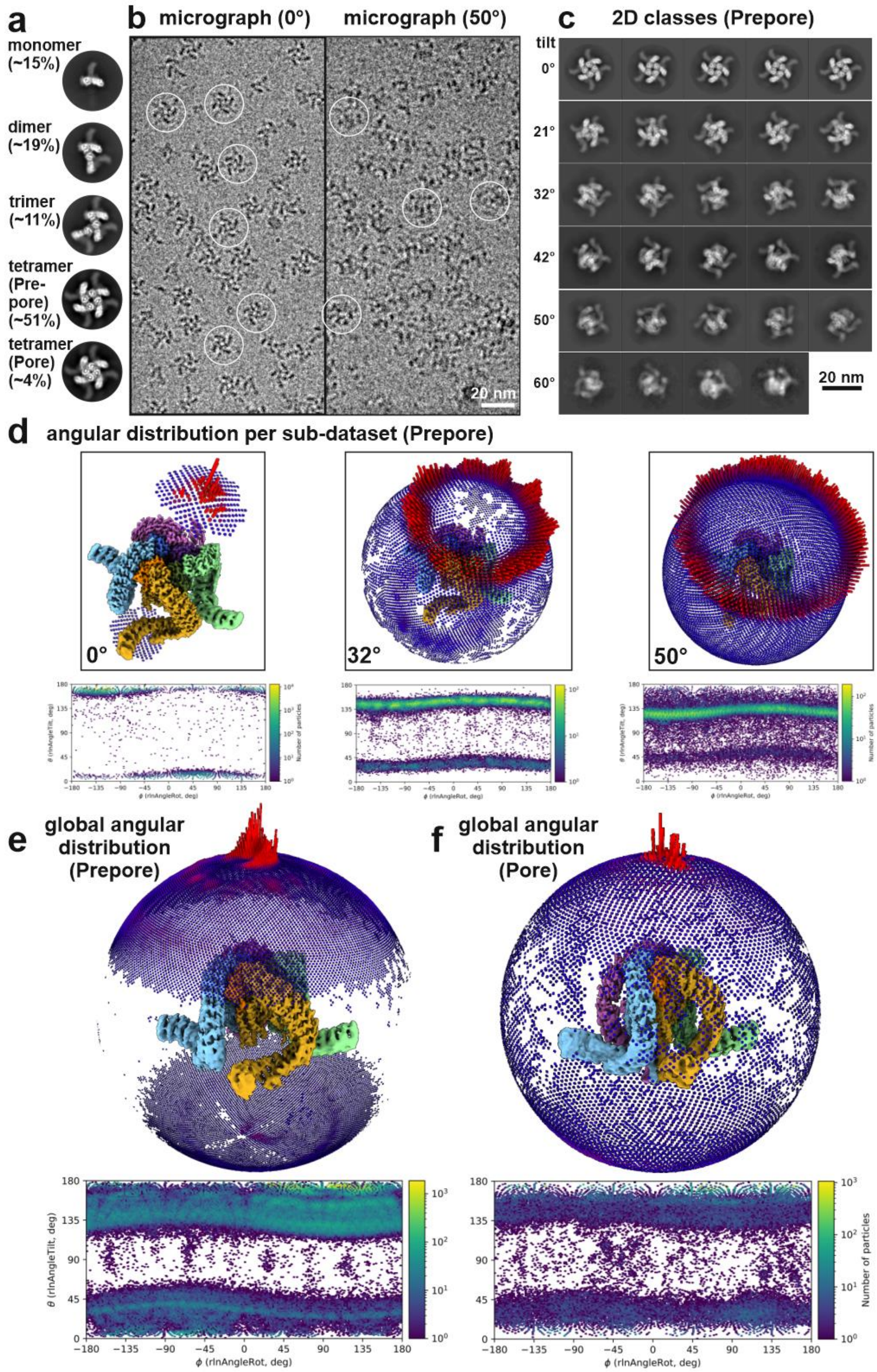

**Supplementary figure 1: Cryo-EM analysis of α-LTX. a** Representative reference-free 2D class averages and distribution of the different oligomeric states from the 0° dataset. Note that the 2D class averages of tetramers show exclusively top-views. **b** Representative cryo-EM micrographs of α-LTX from the untilted and 50° tilted dataset. **c** 2D class averages from datasets collected at 0°, 21°, 32°, 42°, 50°, 60° **d** Distribution of particles in 3D and 2D histogram representation for the 0°, 32° and 50° datasets. **e,f** Particle distributions for the final reconstructions of the prepore and pore state from the merged dataset.

#### Supplementary figure 2

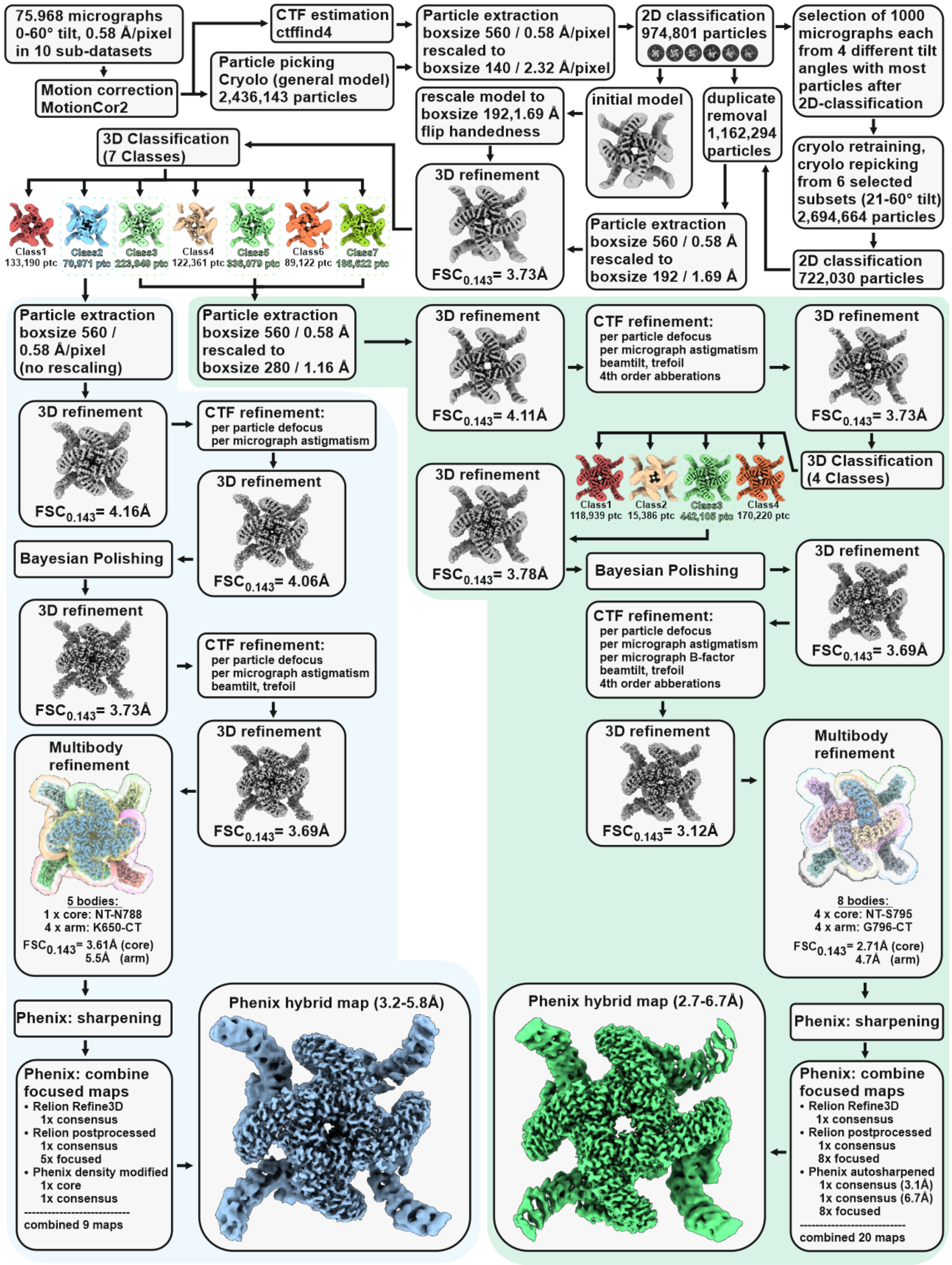


**Supplementary figure 2: Cryo-EM processing workflow for structure determination** of **α-LTX in prepore and pore state.** The flowcharts of α-LTX prepore and pore processing workflows are highlighted with a green and blue background, respectively.

**Supplementary figure 3**

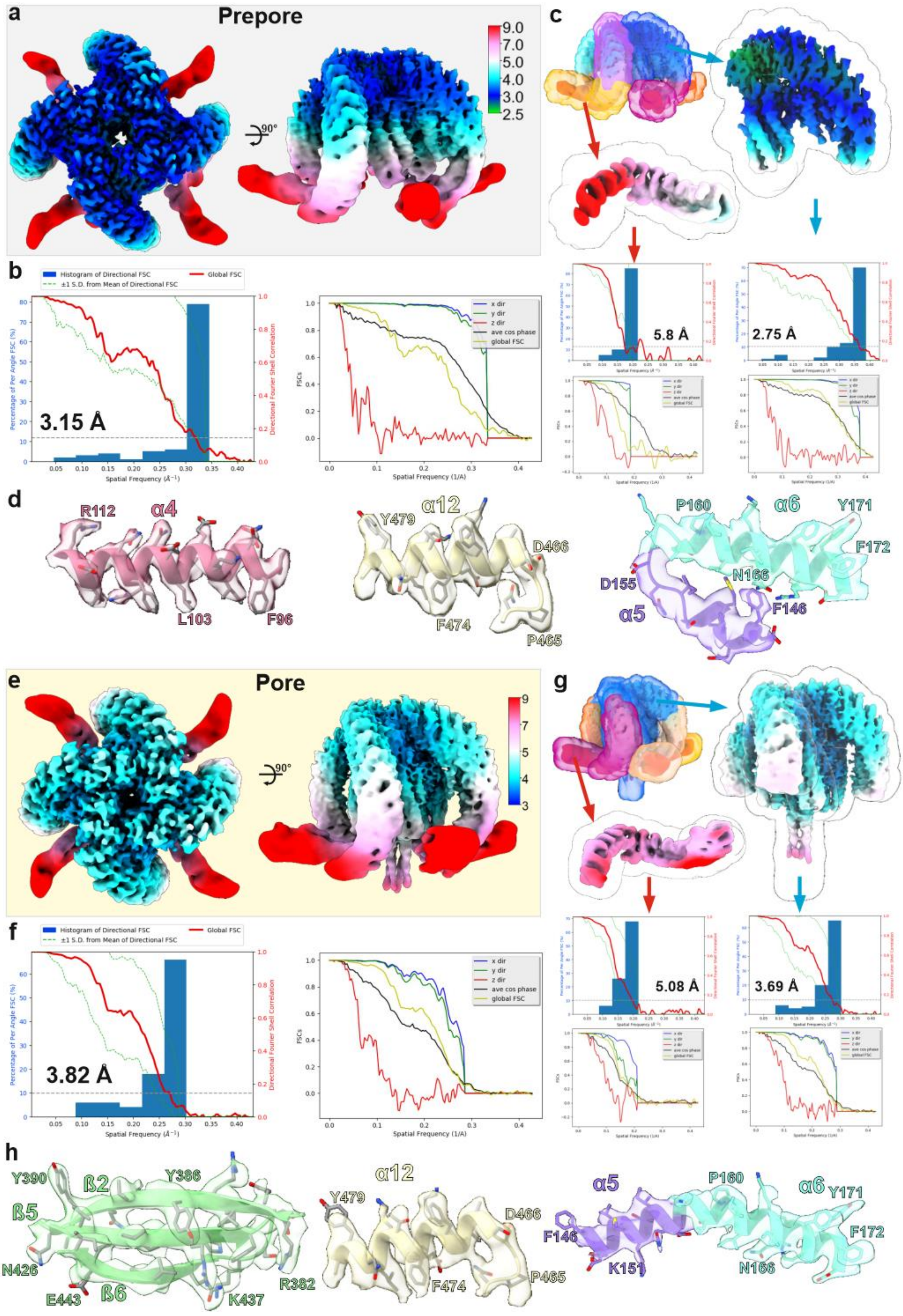

**Supplementary figure 3: Resolution of cryo-EM maps. a,b** Density map of α-LTX prepore prior multi-body refinement colored by local resolution (**a**) and respective 3D-FSC[1] (**b**). **c** 3D masks used for multi-body refinement of the prepore state in RELION4 and the resulting locally refined bodies colored by local resolution, together with their 3D-FSCs. **d** Superposition of segments of the molecular model of the prepore with the composite map for representative regions of the structure with varying local resolution. **e-f** Density map of α-LTX pore prior multi-body refinement colored by local resolution (**e**) and respective 3D-FSC (**f**). **g** 3D masks used for multi-body refinement of the pore state in RELION4 and the resulting locally refined bodies colored by local resolution, together with their 3D-FSCs. **h** Superposition of segments of the molecular model of the pore with the composite map for representative regions of the structure with varying local resolution.

**Supplementary figure 4**

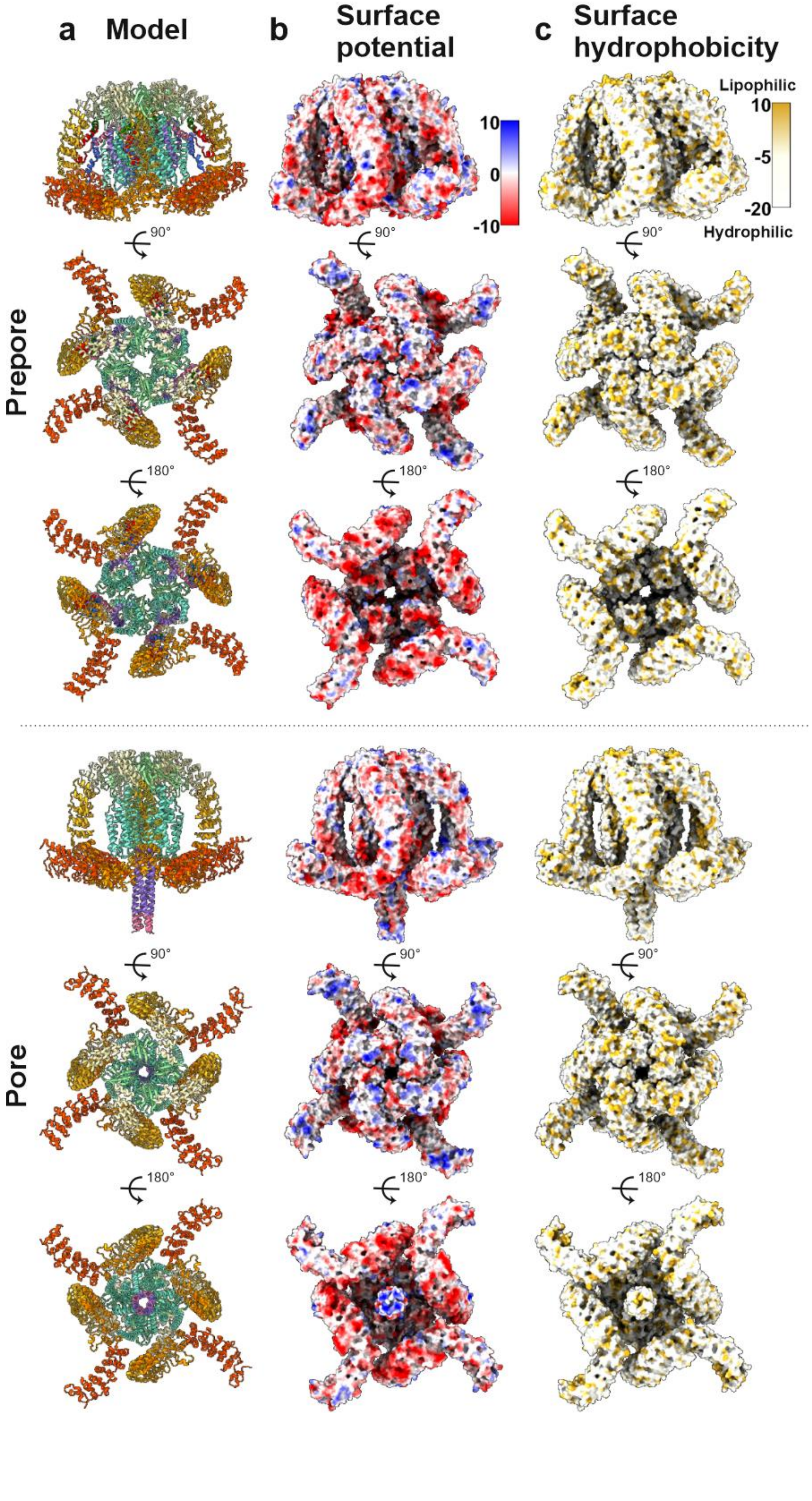

**Supplementary figure 4: Biophysical properties of α-LTX prepore and pore state. a-c** Molecular model (**a**), surface coulombic electrostatic potential at 298K [kcal/(mol·*e*)] (**b**) and Molecular Lipophilicity Potential (MLP) maps (**c**) of the cryo-EM structures in prepore and pore state.

**Supplementary figure 5**

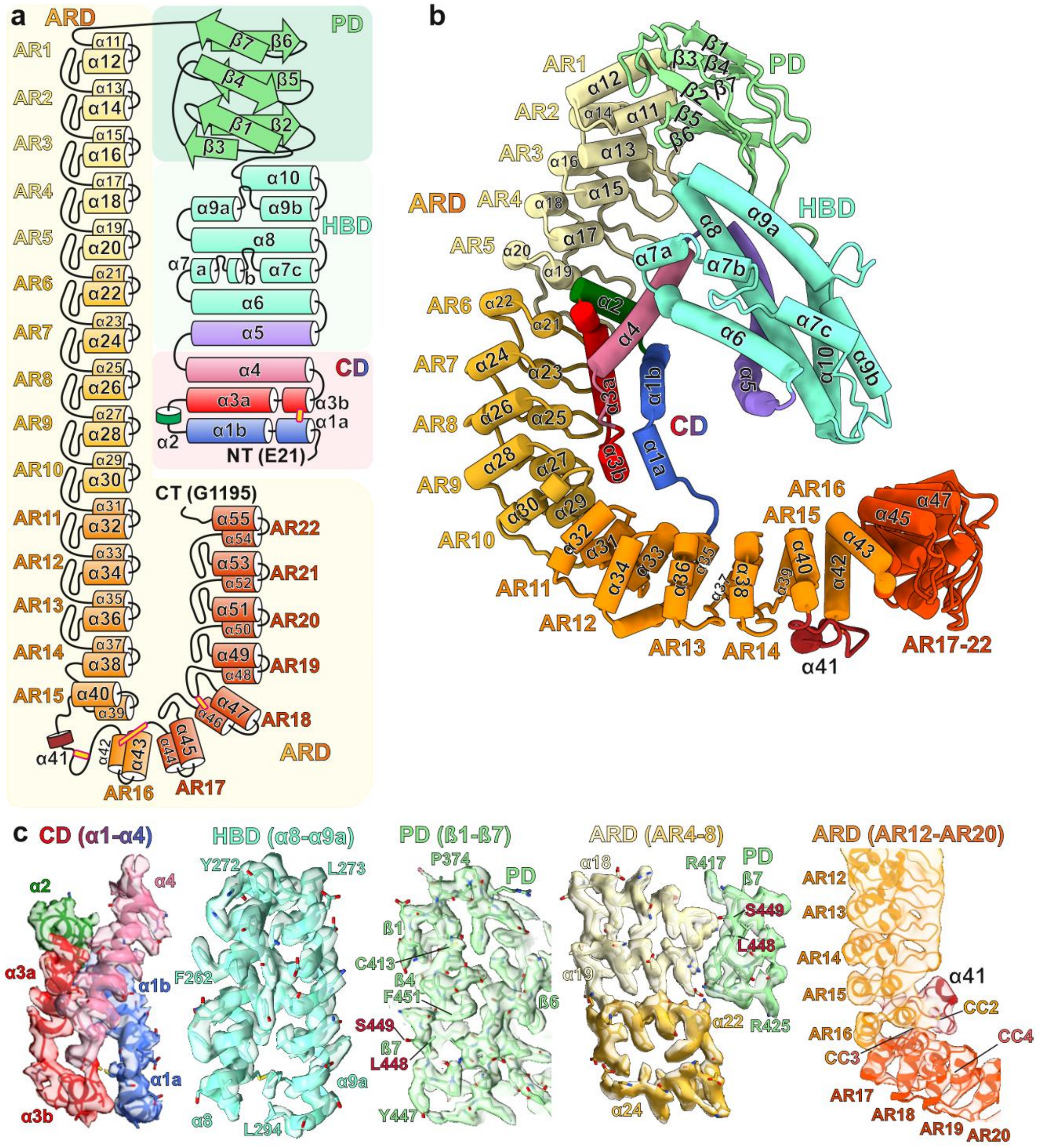

**Supplementary figure 5: Architecture of one subunit of α-LTX tetramer in the prepore state.** **a,b** Topology diagram **(a)** and ribbon representation (**b**) of α-LTX in the prepore state. Connector domain (CD), helical bundle domain (HBD; cyan), plug domain (PD; green) and ankyrin-like repeat domain (ARD; brown spectrum). The nomenclature and color code is used throughout the manuscript. **c** Superposition of segments of the α-LTX prepore cryo-EM density map with the corresponding molecular model for the different domains of the subunit.

**Supplementary figure 6**

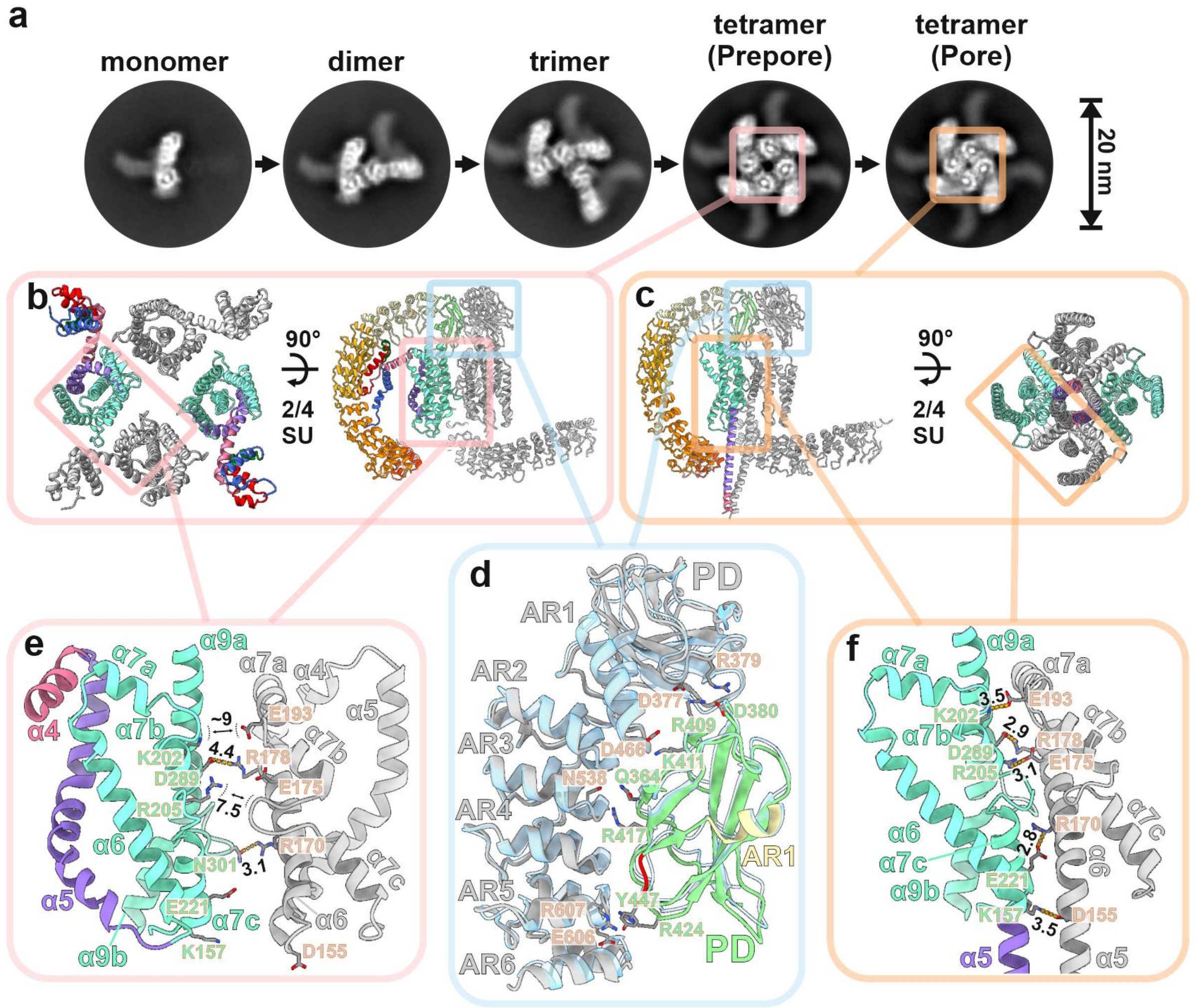


**Supplementary figure 6: Assembly and stabilization of tetrameric α-LTX in the prepore and pore state. a** Stepwise assembly of α-LTX prepore and transition to the pore state. Representative reference-free 2D class averages are shown. **b,c** Top view of the central core of the tetrameric complex (HBD and CD domains) and side view of two clockwise neighboring monomers in α-LTX prepore (**b**) and pore (**c**) state. **d** Interactions between monomers are dominated by contacts between the PD of one subunit with the ARD of the clockwise neighboring monomer. This interface remains stable during the prepore(colored)→pore(grey) transition. **e-f.** Interactions between the HBD domains of adjacent monomers in the prepore (e) and pore (f) state. HBD interactions are rather weak in the prepore state (**e**), but in the pore state neighboring HBD domains move closer together to form a tight interface (**f**).

#### Supplementary figure 7

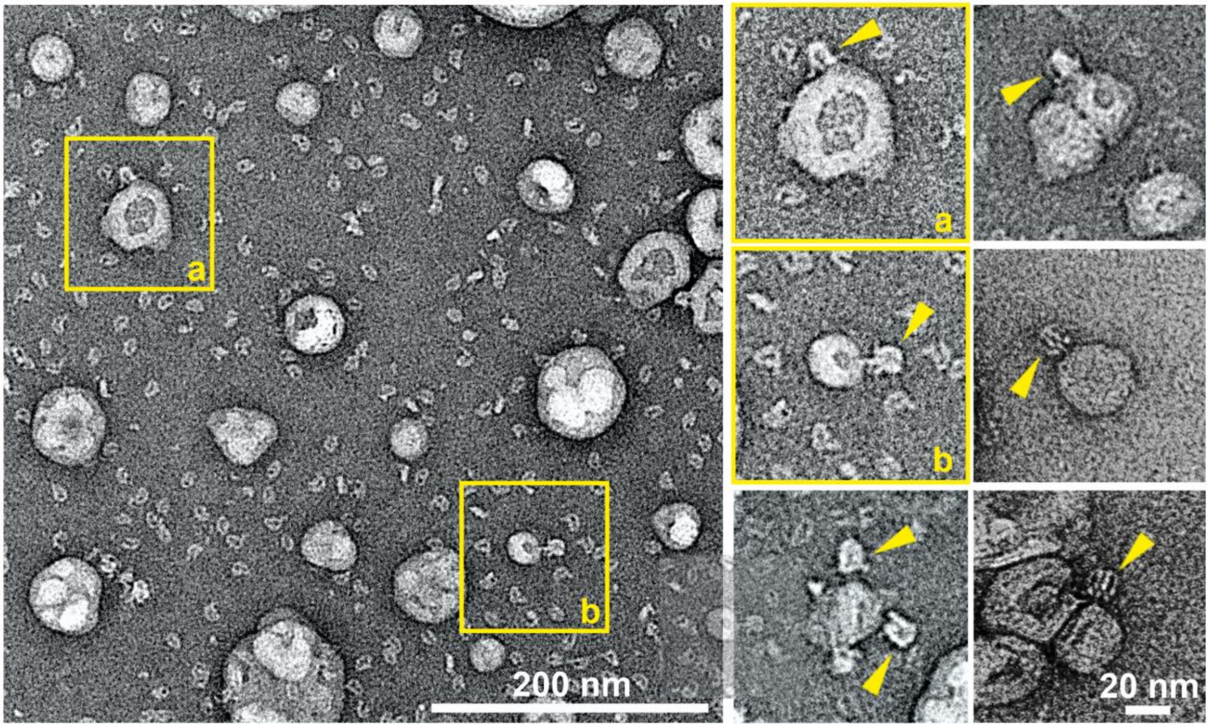


**Supplementary figure 7: α-LTX reconstituted in liposomes.** Representative negative stain EM micrograph (left) and images of individual liposomes (right) with incorporated α-LTX particles (yellow arrows). Particles interacting with the membrane show the characteristic umbrella-like shape of the tetrameric pore. The vast majority of particles are distributed in the background and show the characteristic G-shape of the monomer in the prepore state.

**Supplementary figure 8**

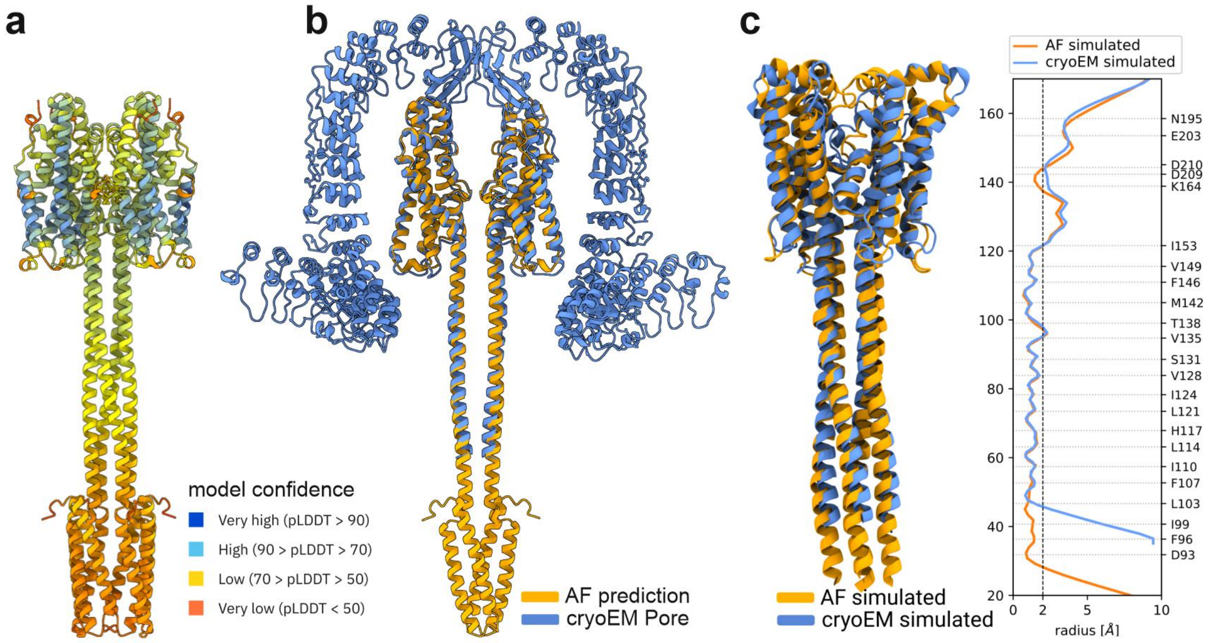


**Supplementary figure 8: Comparison of AlphaFold2 prediction of N-terminal α-LTX central core with the cryo-EM structure of the α-LTX pore. a** AlphaFold2 prediction of a tetrameric N-terminal central core of α-LTX (CD-HBD; residues E21-E360) colored by model confidence **b** Overlay of the AlphaFold2 prediction (orange) with the cryo-EM structure of the α-LTX pore (blue). The RMSD between 221 pruned atom pairs is 0.91 Å. Shown are two opposing subunits. **c** Results of MD simulations of 1 µs, starting from the AlphaFold2 (orange, residues C91 to Y260) and the cryo-EM structure (blue, residues N105 to Y260), respectively. Left: Overlay of both resulting structures. Right: Comparison of the average radius profiles of these structures. One can see the excellent agreement, supporting the close similarity of AlphaFold2 and cryo-EM structures.

#### Supplementary figure 9

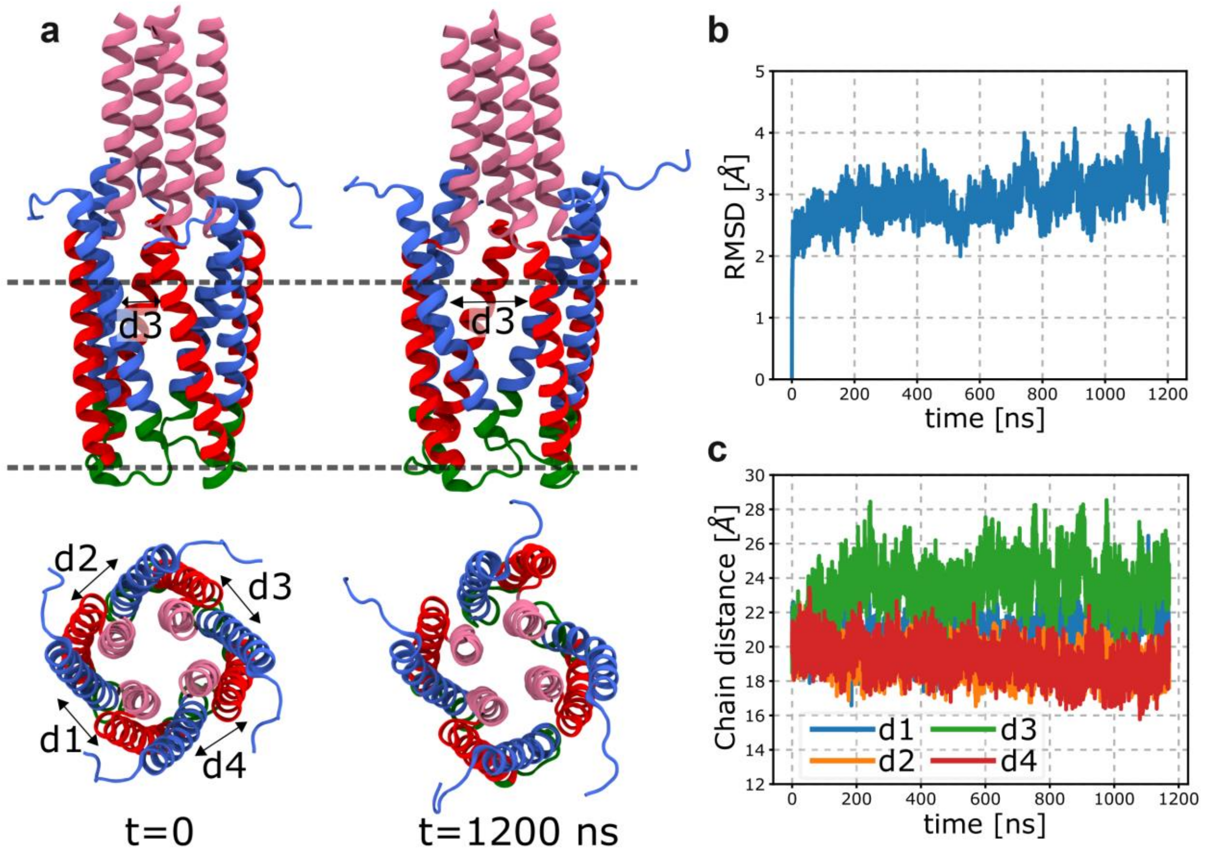


**Supplementary figure 9: MD-simulation of the conformational dynamics and stability of the α-LTX transmembrane pore inside the membrane. a** Structural changes in the TMD of α-LTX predicted by AlphaFold2 inside the membrane during 1200 ns simulation time, obtained via MD simulations (0.15 M $CaCl_2$, no electric field). Chosen are a side and a top view in cartoon representation (lipids not shown). The different orientations help to visualize the residual fluctuations of the TMD. **b,c** RMSD of the protein backbone atoms (**b**) and the average distance between the helix α1 (residues T26-L57) from one subunit to helix α3 (residues G61-G90) of the neighboring subunit (**c**) are shown versus the simulation time in the system with 0.15 M $CaCl_2$. Initially, one of the four distances (d3) increases slightly. Importantly, after about 100 ns, none of the distances increases any more. Thus, the simulations suggest that the overall TMD is stable.

#### Supplementary figure 10

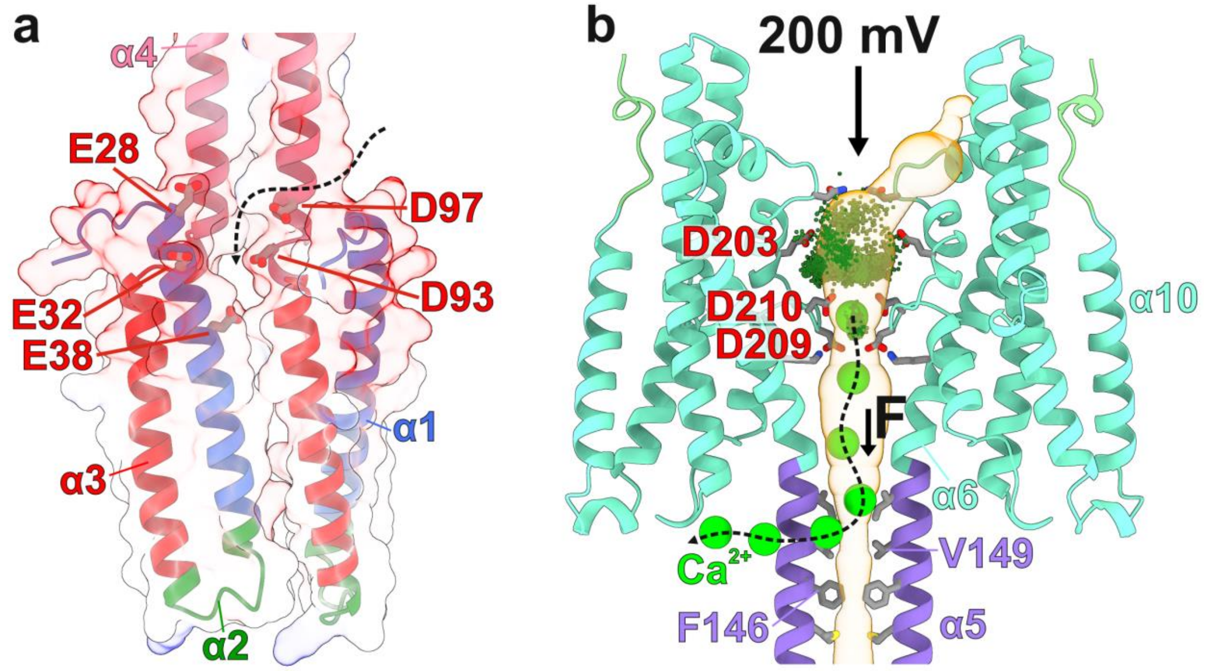


**Supplementary figure 10: Potential cation gates of α-LTX. a** The lateral cation entry site between the distal end of the coiled-coil and the transmembrane domain. Negatively charged residues lining the gate are indicated. The respective MD simulations are shown in Figure 3. **b** The HBD (cyan) and the proximal end of the tetrameric coil (purple) form a continuous channel. The HBD loops form a central cavity with three layers of channel-lining negatively charged residues (E203; D209; D210). MD simulations show that $Ca^{2+}$ ions enter this cavity to bind strongly to the site formed between the second and third layers (D209; D210; diameter 4Å; Figure 3), but do not further enter the coiled-coil stalk. The majority of $Ca^{2+}$ enters the upper opening of the channel, but is less localized, approximately at the level of the first layer (E203; diameter 6Å; Figure 3). The small green dots show the superposition of $Ca^{2+}$ ions from all 1000 frames (step 1 ns) of an MD simulation with an applied electrical potential difference of 200 mV. When an additional pulling force is applied (see methods), $Ca^{2+}$ ions do pass through the strong binding site but then they exit laterally and do not translocate through the stalk. The large green spheres show representative frames of such a translocation event.

**Supplementary figure 11**

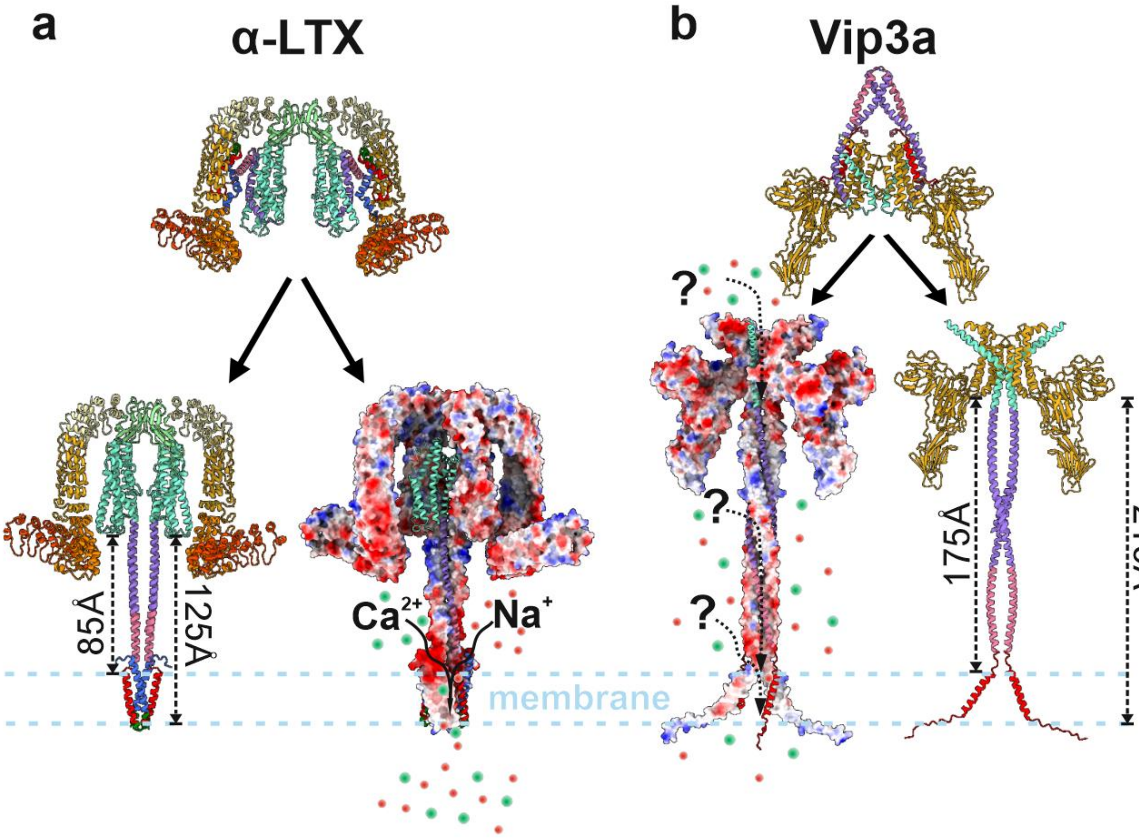


**Supplementary figure 11: Comparison of the pore-forming toxins α-LTX (a) and Vip3a[2] (b)**. Both toxins form channels which are highly cation selective. For each toxin, the prepore state is shown on top and the pore state (upon membrane insertion) at the bottom. Note the characteristic umbrella-like shape of both toxins and also the central tetrameric coiled-coil stalk, required for membrane insertion. The tetrameric assembly of the N-terminus including the potential transmembrane domains was predicted by Alphafold2 in both cases. Surface electrostatics depict the amphipathic properties of the transmembrane domains (TMDs). It should be noted that the prediction of the Vip3a TMD was less stable than for the TMD of α-LTX, and its properties will require further analysis. In contrast to α-LTX, Vip3a does not contain a stabilizing disulfide in this region, suggesting that such an arrangement might contain only one transmembrane helix per monomer and not two like in α-LTX. The cation translocation pathway in Vip3a remains unclear, as does whether the lower end of the stalk provides a lateral entry gate for cations, similar to α-LTX. Further studies are needed to unravel such similarities in the channel-forming mechanisms of these until recently unrelated cation channel-forming toxins.

**Supplementary figures 12:**

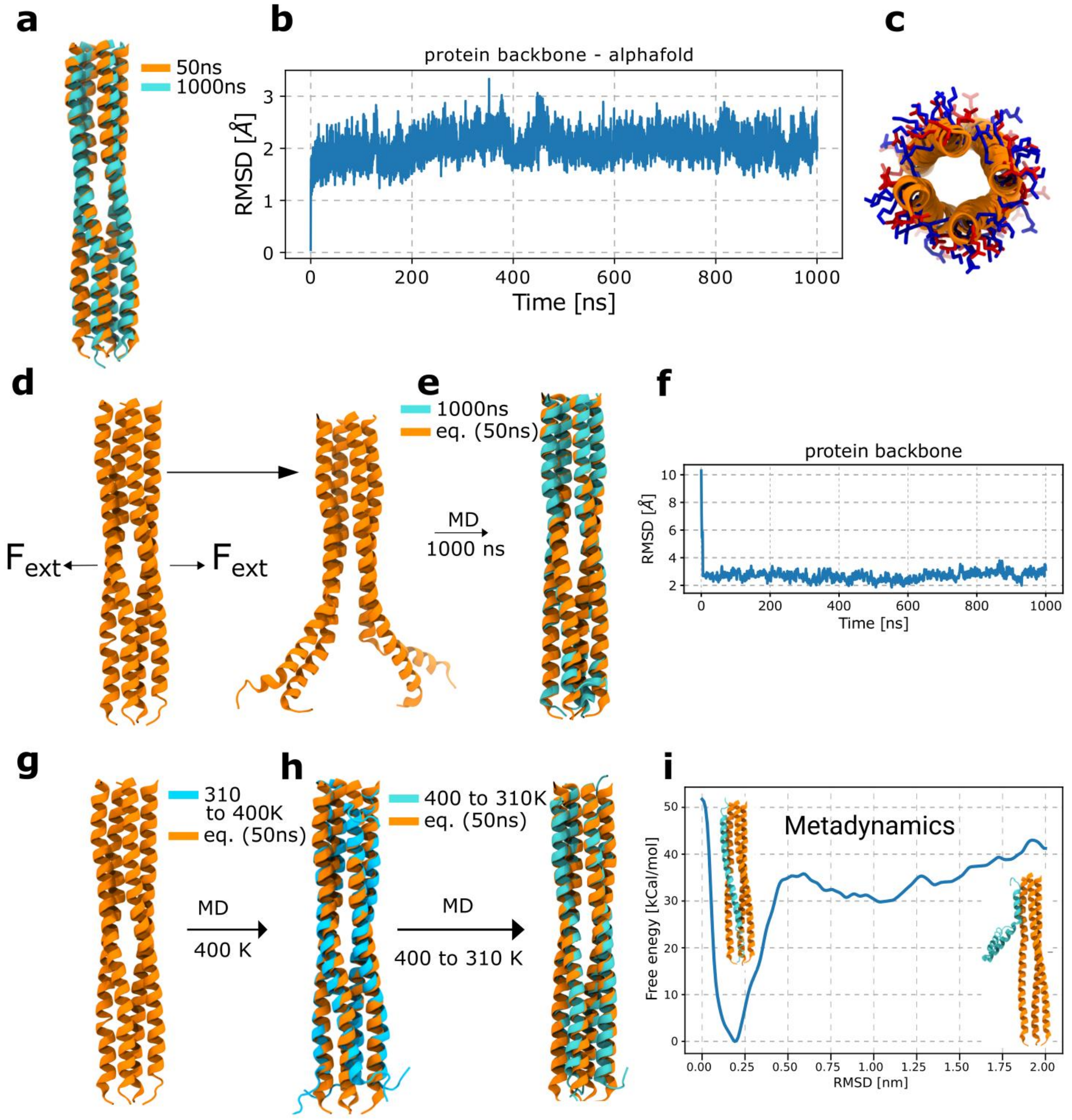


**Supplementary figure 12: MD simulations of the stalk region of the pore structure of the α-LTX predicted by AlphaFold2 to probe its stability. a** Overlay of the cartoon representation of the two structures of the protein from the MD simulations of the stalk (residues C91-D155) in water extracted from the 50 ns (orange) and 1000 ns (cyan) of the simulation, showing that the stalk is quite stable and hardly changes during a microsecond simulation time. **b** The RMSD of the backbone atoms of the stalk as a function of time, which additionally indicates the high stability of the stalk. **c** Top view of the stalk in cartoon representation together with the negatively (red) and positively (blue) residues in sticks representation. The high stability of the stalk is likely due to the specific arrangement of the polar amino acids pointing towards the outside of the pore and the hydrophobic residues inside the pore. **d** Steered MD simulations of the AlphaFold2 structure in which an external force is applied to the center of mass of the four chains to a direction pointing to the outside of the pore. As a consequence, the pore starts to open from the lower part of the stalk. **e** A structure from the steered MD was adopted and a normal MD was performed for 1 µs. The overlay of the structure at the end of this simulation and the equilibrated structure from the normal MD in **a** is represented, showing that the distorted structure due to the steered MD can go back to the initial structure. **f** The RMSD of the protein backbone atoms in the simulations in **e** is shown, denoting that the distorted structure in steered MD abruptly goes to the initial structure and remains stable throughout a microsecond simulation. **g** Simulation of the equilibrated stalk (50 ns) was performed at 400 K, distorting the structure. The overlay of the structure at the end of the simulation (blue) with the equilibrated structure (orange) is shown. **h** Starting with the structure at the end the simulation at 400 K the temperature was gradually decreased to 310 K. The overlay of the structure at the end of this simulation (cyan) and the equilibrated structure (orange) is shown in cartoon representation. Both results shown in **e** and **h** indicate that the stalk structure is indeed close to a deep minimum of the free energy of the system. **i** Free energy profile of stalk formation. The free energy as a function of the reaction coordinate, starting from a combination of three monomers from the tetrameric coiled-coil of the pore state (residues C91-D155, orange cartoon), with one monomer of the prepore structure (cyan), as shown at the right side. The structure at a free energy minimum (at ~0.19 nm) is shown at the left side.

**Supplementary figure 13:**

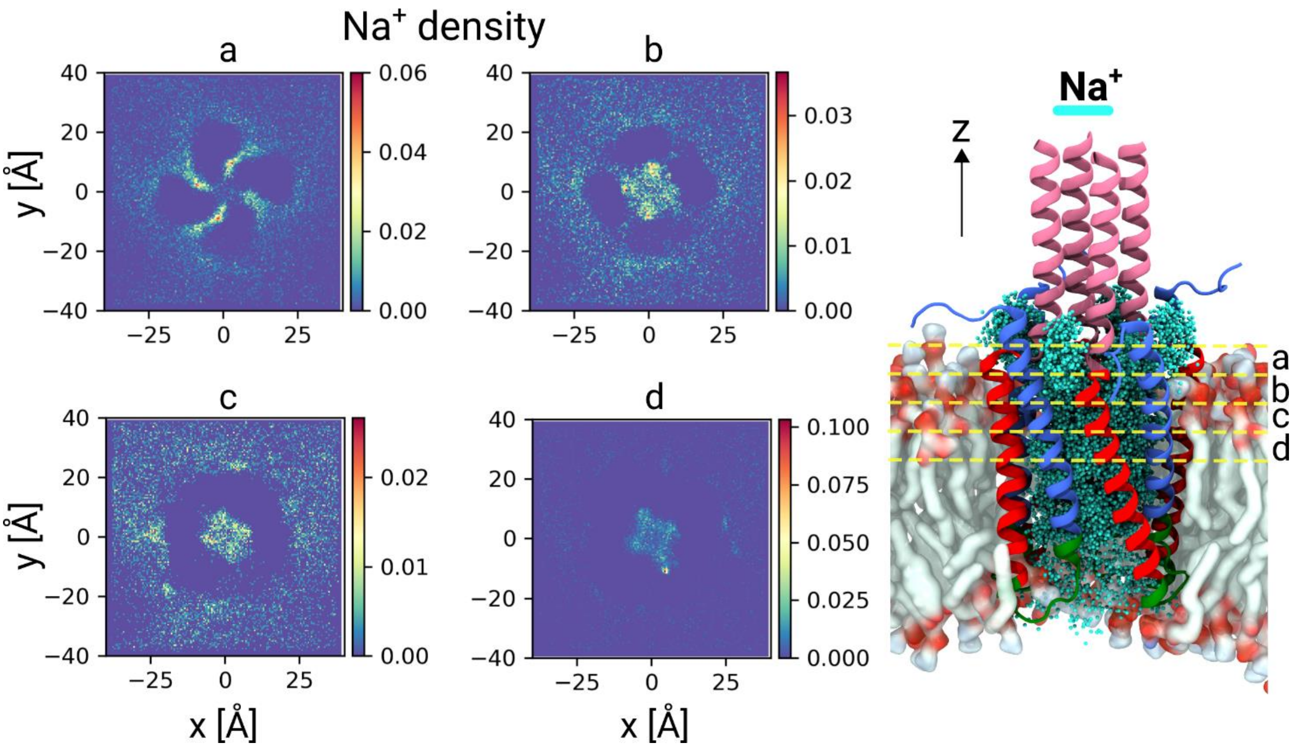


**Supplementary figure 13: Simulations of the membrane part with $Na^+$.** Density maps of $Na^+$ ions from the MD simulation of the TMD of the AlphaFold2 prediction of α-LTX for different cross sections of the protein along the z-axis and width of 5 Å. The corresponding density maps for $Ca^{2+}$ are shown in Figure 3d.

#### Supplementary figure 14

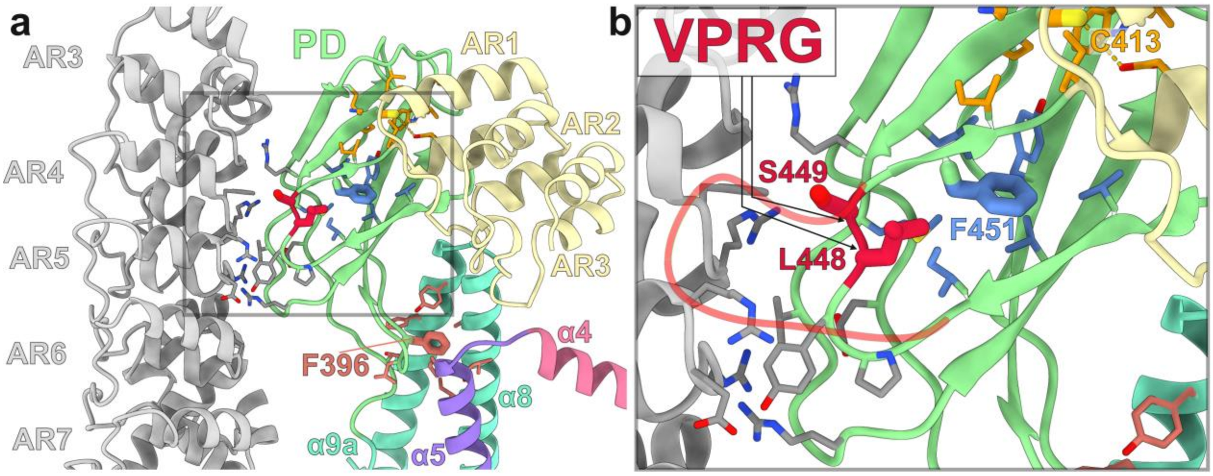


**Supplementary figure 14: Effect of the mutant α-LTX$^{N4C}$ [2]. a** Region of inter- and intrasubunit interactions between the PD and the ARD. For clarity, three hydrophobic pockets with the central atoms C413, F451 and F396 are highlighted in orange, blue and pink, respectively. Residues L448 and S449, between which the VPRG sequence is inserted in the α-LTX$^{N4C}$ mutant[2], are colored in red. **b** A loop created by the VPRG insertion (pale red line) was roughly modeled in coot and superimposed on the WT model to highlight potential clashes with the ARD (gray) of the neighboring subunit, explaining the abolished ability of the α-LTX$^{N4C}$ variant to form tetramers.

#### Supplementary figure 15

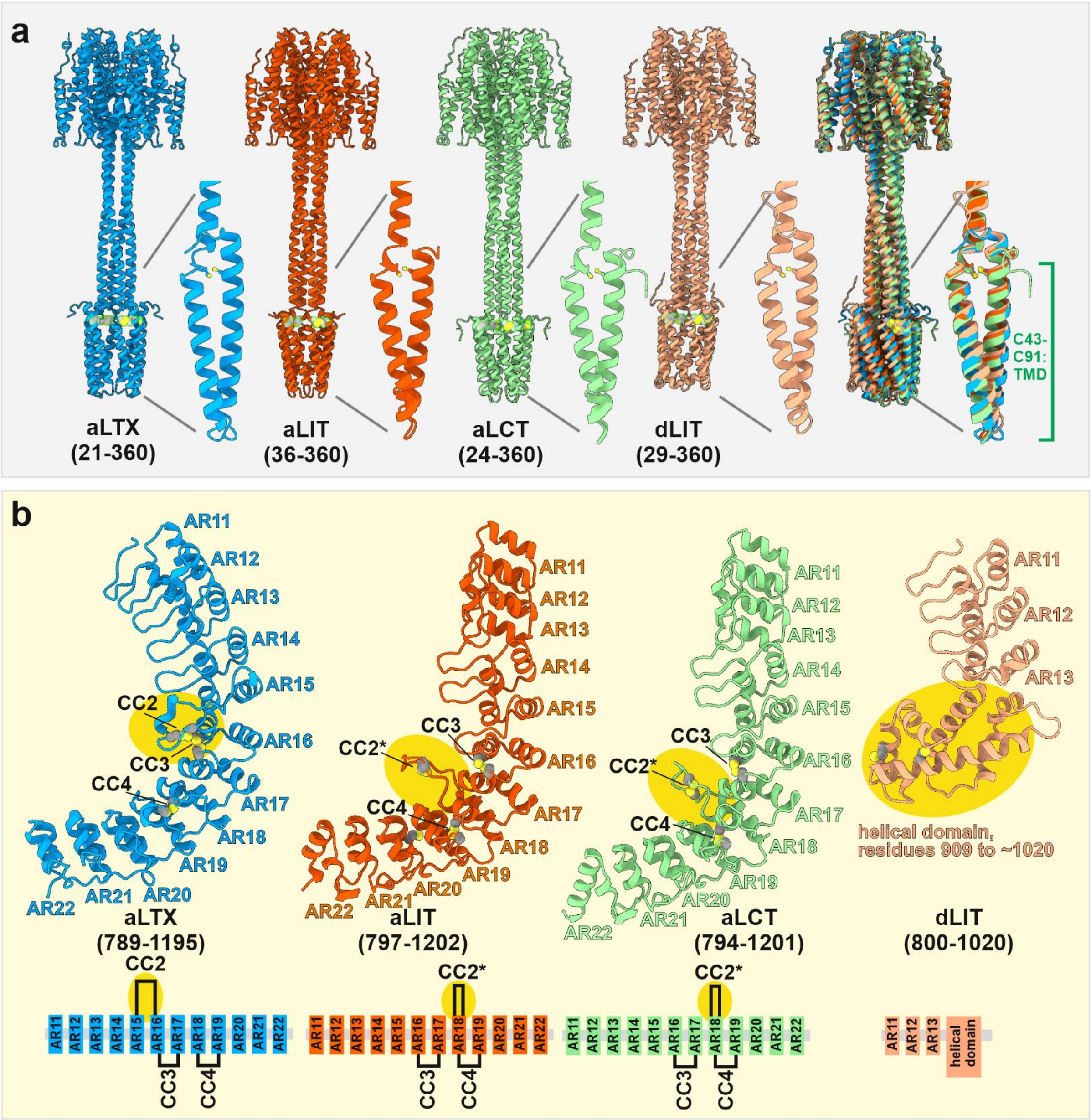


**Supplementary figure 15: Comparison of AlphaFold2 predictions for different latrotoxins (α-LTX (vertebrate-specific), α-LIT, δ-LIT (insect-specific) and α-LCT (crustacean-specific)). a** AlphaFold2 predictions of tetramers of the tetrameric central-core (CD; HBD) of different LTXs. The motif of an N-terminal TMD stabilized by a conserved disulfide bridge (yellow) is predicted in all analyzed latrotoxins. All latrotoxins present a remarkable similarity to α-LTX in this region, with RMSDs of 1.01 Å (α-LIT, 290 pruned atom pairs), 1.08 Å (α-LCT, 243 pruned atom pairs) and 1.03 Å (δ-LIT, 280 pruned atom pairs) **b** AlphaFold2 predictions of monomers of the C-terminal part (ARD) for different latrotoxins. An elongated loop containing a stabilizing disulfide bridge is present between AR15 and AR16 in α-LTX and within AR18 in α-LIT and α-LCT. In contrast, δ-LIT is truncated and ends with a small and highly negatively charged helical domain that contains two cysteines.

#### Supplementary table 1: Cryo-EM Table

| **Data collection** | | |
|---|---|---|
| Microscope | Titan Krios G4<br>(Selectris X, E-CFEG) | |
| Voltage (kV) | 300 | |
| Camera | Falcon 4i | |
| Pixel size (Å) | 0.58 | |
| Tilt angle | Micrographs | Particles |
| 0° | 8,417 | 447,887 |
| 21° | 23,158 | 504,019 |
| 30° | 14,167 | 425,266 |
| 32° | 8,755 | 234,017 |
| 35° | 1,137 | 33,715 |
| 40° | 965 | 20,343 |
| 42° | 11.926 | 320,948 |
| 42° | 2,005 | 74,390 |
| 50° | 4,700 | 359,204 |
| 60° | 738 | 16,354 |
| total number of particles | 2,436,143 | |
| Number of frames | 918-1225 | |
| Number of fractions | 49-51 | |
| Total electron dose ($e^-$/Å$^2$) | 50 | |
| Defocus range (µm) | -0.3 – -1.7 | |
| **Atomic model composition** | **Prepore** | **Pore** |
| Chains | 4 | 4 |
| Symmetry imposed | C1 | C1 |
| Non-hydrogen (protein) atoms | 36,980 | 34,612 |
| Residues | 4,700 | 4,364 |
| particle substack | 442,105 | 70,971 |
| Ligand atoms | - | - |
| **Refinement (Phenix)** | | |
| RMSD bond | 0.005 | 0.007 |
| RMSD angle | 0.753 | 0.847 |
| Model to map fit, CC mask | 0.76 (0.75) | 0.67 (0.66) |
| Model to map fit, CC box | 0.60 (0.87) | 0.71 (0.76) |
| Resolution (FSC@0.143, Å) | 2.71 (3.15) | 3.61 (3.69) |
| B-factor (mean, Å$^2$) | 117.45 | 121.68 |
| **Validation** | | |
| Clashscore | 13.64 | 15.79 |
| Ramachandran outliers (%) | 0 | 0.07 |
| Ramachandran favoured (%) | 95.89 | 96.05 |
| Molprobity score | 2.48 | 1.97 |
| EMRinger score | 1.50 (1.38) | 1.41 (0.57) |

**Supplementary table 1: Cryo-EM data collection and refinement statistics of α-LTX.** The refinement in Phenix was done against hybrid maps obtained by Phenix.combine_focussed_maps after multi-body refinement (EMDB-ID XXX); Supplementary figure 2). The presented refinement statistics are for the hybrid maps (resolution is the reported resolution from multi-body refinement), and in brackets for the same model against the original, not locally refined nor combined maps after 3D-refinement and sharpening in RELION4.

#### Supplementary table 2: Number of TMD permeation events

| $Na^{+}/Ca^{2+}/La^{3+}$ | $Na^{+}$ | $Na^{+}$ with EF | $Ca^{2+}$ | $Ca^{2+}$ with EF | $La^{3+}$ | $La^{3+}$ with EF |
|---|---|---|---|---|---|---|
| ext. to int. | 51 | 136 | 6 | 61 | 0 | 0 |
| int. to ext. | 43 | 19 | 4 | 1 | 0 | 0 |

**Supplementary table 2:** Number of permeation events of the cations across the protein membrane part during the production simulation of 1200 ns, starting from the AlphaFold2 structure, for different ions and with and without the applied electric field corresponding to an electrical potential difference of 100 mV. As expected, without an electric field the number of transitions is, within statistical uncertainties, identical in both directions. Application of an electric field induces a directional bias due to the additional gradient in potential energy which is particularly pronounced for the divalent cation.

#### Supplementary table 3: Average time duration of TMD permeation events

| $Na^{+}/Ca^{2+}/La^{3+}$ | $Na^{+}$ | $Na^{+}$ with EF | $Ca^{2+}$ | $Ca^{2+}$ with EF | $La^{3+}$ | $La^{3+}$ with EF |
|---|---|---|---|---|---|---|
| ext. to int. | 25 | 19 | 170 | 76 | - | - |
| int. to ext. | 31 | 18 | 251 | 93 | - | - |

**Supplementary table 3.** Average time duration (in ns) of the permeation events of cations across the membrane part, as listed in Supplementary Table2. As expected from transition state theory, despite application of an electric field the transition times remain similar in both directions but become shorter. The transition times for the divalent cation is approx. 3 times slower as compared to the monovalent cation.

#### Supplementary table 4: Interfaces between α-LTX subunits

| Prepore | Inter-molecular interfaces | | |
|---|---|---|---|
| **Fragment A residues** | **Fragment B residues** | **$\Delta^iG$ (A-B) [kcal/mol]** | **Interface area (A-B) [Å$^2$]** |
| I454-G1195 | Q351-D453 | -3.3 ± 0.4 | 692.4 ± 12.9 |
| A156-K350 | E21-C91 | -0.7 ± 0.6 | 30.9 ± 22.2 |
| **Prepore** | **Intra-molecular interfaces** | | |
| **Fragment A residues** | **Fragment B residues** | **$\Delta^iG$ (A-B) [kcal/mol]** | **Interface area (A-B) [Å$^2$]** |
| A156-K350 | S92-D155 | -21.4 ± 1.7 | 1595.0 ± 46.2 |
| I454-G1195 | E21-C91 | -25.0 ± 1.4 | 1447.3 ± 59.5 |
| S92-D155 | E21-C91 | -15.8 ± 0.4 | 735.7 ± 9.3 |
| I454-G1195 | S92-D155 | -3.7 ± 0.7 | 310.1 ± 5.1 |
| Q351-D453 | S92-D155 | -5.2 ± 0.1 | 204.5 ± 2.2 |

| Pore | Inter-molecular interfaces | | |
|---|---|---|---|
| **Fragment A residues** | **Fragment B residues** | **$\Delta^iG$ (A-B) [kcal/mol]** | **Interface area (A-B) [Å$^2$]** |
| S92-D155 | S92-D155 | -21.5 ± 0.8 | 1297.2 ± 26.0 |
| S92-D155 *(diagonal contact)* | S92-D155 *(diagonal contact)* | -5.10 ± 0.2 | 166.8 ± 3.0 |
| A156-K350 | A156-K350 | -4.1 ± 0.5 | 856.9 ± 35.7 |
| I454-G1195 | Q351-D453 | -2.2 ± 0.8 | 714.3 ± 21.5 |
| E21-C91 | E21-C91 | -4.7 ± 1.8 | 429.4 ± 72.1 |
| A156-K350 | S92-D155 | -0.7 ± 0.04 | 23.4 ± 1.6 |

**Supplementary table 4: Interfaces between α-LTX subunits in the tetrameric assembly.** Each of the four α-LTX monomers were segmented into five fragments, and mutual interactions between fragments A and B as indicated were analyzed by the Pisa server[3]. Residues E21-C91 correspond to helices α1-α3 that form the transmembrane domain in the pore, residues S92-D155 form the tetrameric coiled-coil stalk in the pore state, residues A156-K350 corresponds to the HBD excluding helix α5 which is part of the stalk in the pore state, residues Q351-D453 correspond to the PD, and I454-G1195 to the ARD. Only interactions with significant relevance for stability are shown. Clockwise intermolecular interactions within the tetramer are averaged; only within the stalk region, interactions of diagonally opposing monomers are also contributing significantly to complex stability. Interactions between residues E21-D155, which form the stalk and transmembrane domain in the pore state, contribute by far strongest for the stability of the tetrameric α-LTX pore. Since intermolecular interactions of these residues are missing in the prepore state, the tetrameric prepore is much weaker than the pore assembly. However, in the prepore state, residues E21-D155 are involved in intramolecular interactions which sum up to slightly higher solvation free energies ($\Delta^iG$) than their intermolecular interactions in the pore state, indicating that despite the high solvation free energy of the formed coiled-coil stalk, the prepore to pore transition is not an exothermic process.

**Supplementary video 1** Cryo-EM map of α-LTX prepore and the corresponding molecular model.

**Supplementary video 2** Cryo-EM map of α-LTX pore and the corresponding molecular model.

**Supplementary video 3** Conformational variability of the prepore state of α-LTX. Shown are morphs between cryo-EM 3D classes of the α-LTX prepore state (Supplementary figure 2).

**Supplementary video 4** Time evolution of the TMD region during 1200 ns, as obtained from the MD simulations. The initial and final configurations of this video are shown in Supplementary figure 9.

**Supplementary video 5** Simulations of the TMD with $Na^+$ and $Ca^{2+}$ ions for an applied electric field corresponding to an electric potential difference of 100 mV. Shown are the protein and the different ions for a time duration of approx. 150 ns. The time evolution is identical for both ions, highlighting the faster translocation of $Na^+$ through the channel. The lipids are hidden for better clarity.

**Supplementary video 6 $Ca^{2+}$-binding stabilizes the tetrameric prepore in a narrow conformation and thereby assists pore formation.** Morphs between different cryo-EM 3D classes of the α-LTX prepore and pore state. Note the "breathing motions" of the central channel in the prepore state. Narrowing of the channel and its stabilization in this conformation by $Ca^{2+}$ assists the pore formation events.

**Supplementary video 7** The video shows a simplified model of α-LTX prepore→pore transition, membrane penetration, channel formation and $Ca^{2+}$ translocation obtained after morphing between the structures in the prepore and the pore state. It should be noted that structures of possible intermediate states, as shown in this animation, are not yet available.

**Supplementary video 8** α-LTX prepore→pore transition (for description see Supplementary video 7), as seen from the bottom- and top-view.

#### Materials and Methods

##### Protein purification

The α-LTX sample used in the study was purchased from Alomone Labs (Cat. LSP-130). It was isolated from *Latrodectus tredecimguttatus* (black widow spider) venom by modifying a previously reported protocol[4,5]. The lyophilized protein was dissolved in sample buffer containing 25 mM Tris-HCl pH 8.0, 150 mM NaCl, 0.1 mM $CaCl_2$, and 0.2 mM $MgCl_2$ and further purified by size exclusion chromatography on a Superdex 200 increase 5/150 column (Cytiva) equilibrated in sample buffer. The eluates were fractionated and confirmed by SDS-PAGE and negative stain electron microscopy.

##### Liposome preparation and binding assay

For liposome preparation, 10 mg 1-palmitoyl-2-oleoyl-sn-glycero-3-phosphocholine (POPC) lipids (Avanti Polar Lipids) were dissolved in a 1:1 methanol/chloroform mixture and lyophilized as a thin lipid film. The film was then rehydrated with 1 ml buffer-detergent mix, containing 1% Triton-X 100 in sample buffer. After dissolving, the mixture was incubated in a glass vial with 10% (w/v) Bio Rad Bio-Beads SM-2 Adsorbent Media (Cat No. 152-3920) for 30 minutes at 4 degrees. Subsequently, Bio-Beads were added to a total of 30% (w/v), and the mixture was incubated overnight at 4°C to remove all traces of detergent. As an alternative, we also prepared liposomes by the extrusion method, without using detergents: a suspension of 10 mg/ml POPC was homogenized by passing 21 times through a polycarbonate membrane with a 0.2 µm pore size in a mini extruder (Avanti Polar Lipids).

For the binding assay, the supernatant was diluted with sample buffer complemented with 5 mM $CaCl_2$ and 0.5 M KCl and incubated with α-LTX at final concentrations of 0.1 mg/ml liposomes and 0.02 mg/ml α-LTX for four hours at 310 K. The membrane-incorporation of α-LTX was then analyzed by negative stain electron microscopy.

##### Negative stain electron microscopy (EM) analysis

For negative stain EM sample preparation, 4 µl of the protein sample, diluted to a concentration of 0.005-0.02 mg/ml, was applied to a glow-discharged carbon-coated copper grid, and incubated for 2 min at room temperature. The excess protein solution was removed by blotting with Whatman (grade 4-5) filter paper, followed by washing with 2x10 µl deionized water or sample buffer, and 2x10 µl 0.75% uranylformate solution. The final uranylformate droplet was incubated for 45-60 seconds before blotting and air drying of the grid.

Image acquisition was performed using a Talos L120C G2 TEM operating at an acceleration voltage of 120 kV. Datasets were acquired with a 4k × 4k CETA-F scintillator camera at a defocus of −1 µm and a pixel size of 1.2 Å/px.

##### Sample vitrification and cryo-EM data acquisition

For preparing the grids for cryo-EM, 4 µl of α-LTX sample at a concentration of 0.5 mg/ml were applied onto a freshly glow-discharged UltrAuFoil R2/1 holey gold grid (Quantifoil). After removal of excess liquid, the sample was vitrified in liquid ethane using a Vitrobot II automatic plunge-freezer (Thermo Fisher Scientific).

Datasets were acquired with a 300 kV Titan Krios G4 microscope (Thermo Fisher Scientific) equipped with an E-CFEG, a Selectris X energy filter and a Falcon 4i direct electron detector operated by the software EPU (Thermo Fisher Scientific). A total of ~90.000 micrographs in 10 sub-datasets at different stage tilt angles from 0 to 60 degree were collected in Electron Event Representation mode (EER) at a nominal magnification of 215k, corresponding to a pixel size of 0.58 Å/px. The majority of micrographs were collected at defocus of -0.3 to -1.7 µm. The Selectris X energy filter was used for zero-loss filtration with an energy width of 10 eV. A total dose of ~50 $e^-/Å^2$ (estimated shortly after freshly flashing the cold-FEG) was aimed for by adjusting the exposure time of each sub-dataset to an appropriate value (between 3 and 4 seconds per micrograph). The details of dataset collection are summarized in Supplementary table 1. Micrographs in which no particles were picked during image processing (see below) are not shown in the statistics.

#### Image processing and 3D reconstruction

EER movies were motion corrected in RELION4[6] using its own Motioncorr2[7]-like algorithm. In a first step, 2,436,143 particles were picked from the motion-corrected micrographs by crYOLO[8] based on a generalized picking model and extracted with a final window size of 560 × 560 pixels. CTF estimation was performed using CTFFIND4[9]. Based on the estimated defocus values and resolution limits, 75,968 good micrographs were selected for further processing.

An initial 2D classification was performed in RELION4 at a pixel size of 2.32 Å/px. Classes displaying incomplete tetramers, and contaminants were removed, and the remaining 974,801 particles were used to generate an initial 3D model. From a range of four different tilt angles, 1000 micrographs (for each tilt-angle) with most particles after 2D classification were used to manually train crYOLO for an optimized picking model. Another 2,694,664 particles were picked with this model from the datasets with 21°-60° tilt and also classified in 2D. After combining the best particles with the first set of 2D classified particles and removal of duplicates, 1,162,294 particles were reextracted at 1.69 Å/px. A 3D Refinement in RELION4 reached 3.73 Å resolution. In a subsequent 3D classification into 7 classes, one class clearly showed a largely different (Pore) conformation than the other (Prepore) classes. We used this pore class with 70,971 particles and the three best prepore classes with a combined amount of 746,650 particles for further processing of the pore and the prepore state, respectively. For generating the Supplementary video 3 and Supplementary video 6, particles from all fully assembled prepore classes were used to represent a more complete conformational landscape of prepore motions (classes 1,3,5,6 and 7 with a total of 968,962 particles; Supplementary figure 2).

The pore state was further processed at full pixel size of 0.58 Å/px. A first 3D refinement reached 4.16 Å resolution, and the 3D map was improved to 3.69 Å resolution by one round of CTF and aberration refinement, Bayesian polishing, and another round of CTF and aberration refinement, each followed by 3D auto-refine in RELION4. The obtained map had a strong resolution gradient and presented strong heterogeneity, particularly in the C-terminal part of the ARD regions. This was accounted for by a final multi-body refinement with one body for each C-terminal half of the ARDs and one body for the central part of the complex, which improved the resolution in the center to ~3.2 Å. To maximize map interpretability, we also generated sharpened maps in RELION4 and density modified maps in Phenix[10]. Since each map provided best interpretability in a different region of the structure, we combined a total of 9 maps with Phenix.combine_focused_maps. This approach analyzes the map-model correlation of multiple maps to identify and superpose the best parts of each map to create a composite map[10]. We used this map together with the individual maps for model building and map interpretation (Supplementary figure 2).

The prepore state was processed using a similar approach, but at a pixel size of 1.16 Å/px and with the difference that after a first round of CTF and aberration refinement, a second 3D classification into 4 classes was used to further remove conformational heterogeneity. A class containing 442,105 particles displaying high-resolution features up to 3.78 Å was selected for subsequent Bayesian polishing and another round of CTF and aberration refinement, each followed by 3D auto-refine in RELION 4. This improved the resolution to 3.12 Å. A final multi-body refinement with one body for each C-terminal half of the ARDs and one body for each N-terminal part improved the core resolution to ~2.71 Å, and the C-termini also became more interpretable. Like for the pore state, we generated a combined focused map using Phenix, here with a total of 20 individual maps to generate an optimized composite map.

#### AlphaFold2 structure prediction

A prediction of the monomeric mature α-LTX (UniprotKB ID: P23631 (LATA_LATTR)) was performed using AlphaFold2[11]. It resembled our previously solved structures of α-LCT and δ-LIT (PDBIDs: 7PTX and 7PTY, respectively), with the same domain architecture and a similar G-shaped arrangement[12]. The N-terminal CD domain had a low confidence in the prediction, and assumed a conformation that is not consistent with our cryo-EM prepore structure. Attempts to predict the full tetrameric assembly using the "multimer" mode of AlphaFold2 were not successful, but a prediction of residues E21-E360, corresponding to the CD and HBD, assumed a conformation in which the CD domain and helix α5 of the HBD form an elongated tetrameric coiled-coil stalk. This conformation largely differs from the prediction of the monomer, but instead is consistent with the pore state as identified by cryo-EM and negative stain EM. For all AlphaFold2 predictions, five models were generated and the model with the highest prediction score was used for the further analysis.

#### Model building and validation

The AlphaFold2 prediction of monomeric α-LTX was used as an initial model for building the cryo-EM prepore state. Initially, the model was copied to the four subunits of the experimental tetramer and relaxed into the density by fragmented rigid-body fitting and subsequent real-space refinement in Phenix[10,13]. For the regions where the predicted model did not fit well (particularly the connector domain), the model was rebuilt from scratch using the model editing software COOT[14,15]. The resulting model was further refined using a combination of COOT and Phenix. The model quality was evaluated by the validation tools implemented in Phenix[16] and the wwPDB validation server[17]. Multiple rounds of the above adjustments were performed until the model sufficiently described the experimental map.

An initial model of the pore state was created by combining residues N105-K350 from the AlphaFold2 prediction of the tetrameric N-terminus with the missing C-terminal part from the monomer prediction. Like for the prepore state, fragmented rigid-body fitting and real-space refinement in Phenix followed by multiple rounds of model refinement with COOT and Phenix were used to create the pore model.

#### Visualization and analysis of cryoEM maps and models

Visualization, analysis and figure preparation was done with ChimeraX (UCSF)[18,19]. Local resolution gradients within a map were calculated with RELION4 and visualized with ChimeraX. 3D angular distribution plots were generated in Relion4. 2D histograms of the angular distribution were generated using angdist[20]. The 3D Fourier shell correlation of cryo-EM maps was calculated using the remote 3DFSC processing server[1] . Interfaces within α-LTX tetramers and their solvation free energies were analyzed by the Pisa server[3].

#### MD simulations: system preparation

Starting from the AlphaFold2 prediction, we modeled, first, the tetrameric coiled-coil domain, namely the stalk (residues C91 to D155) and, second, a part of the HBD region and the tetrameric coiled-coil region (residues C91 to Y260). The proteins were then solvated with water and ions using the CHARMM-GUI solvation builder[21]. For the first case 0.15 M NaCl was used, in the second case an additional simulation with 0.15 M $CaCl_2$ was performed. A similar procedure was followed to model the cryo-EM structure of the pore state (residues N105 to Y260) solvated with water and 0.15 M NaCl.

To model the membrane protein part, the residues E21 to S116 were inserted in a 7:2:1 POPC:CHOL:POPS lipid mixture, corresponding to a synaptic membrane, which were then solvated with water and ions, using CHARMM-GUI membrane builder[21]. Three systems were simulated with three different ion concentrations of 0.15 M NaCl, $CaCl_2$ and $LaCl_3$.

#### MD simulations

All MD simulations were performed using the 2019.6 version of GROMACS[22,23]. The TIP3P water model was used to describe the water molecules[24]. Periodic boundary conditions were applied in all directions. The simulation sizes were large enough to avoid protein-protein interactions via periodic images. Long-range electrostatic interactions were treated with the use of the particle mesh Ewald method[25], with a cutoff distance of 1.2 nm and a compressibility value of $4.5 \times 10^{-5}$. The van der Waals (vdW) interactions were treated using cut-off schemes with a cutoff distance of 1.2 nm, which are smoothly truncated between 1.0 and 1.2 nm. The constant pressure was maintained at 1 bar by coupling the system to the Berendsen barostat[26] in equilibration and Parrinello-Rahman[27] barostat in production simulations, using the semi-isotropic pressure scheme for the membrane-protein system. The temperature was controlled at 310 K for all systems otherwise stated by coupling the system to the Nosé-Hoover thermostat[28]. The LINCS algorithm was utilized to constrain the bonds[29]. All systems were first minimized and subsequently equilibrated using initially the NVT (500 ps) and then the NPT (16 ns) protocol in multiple steps. During the course of equilibration, restraints (1000 kJ $mol^{-1}$ $nm^{-2}$) were applied on the protein so that the membrane and solvent are equilibrated around the protein. The production simulations were performed for 1200 ns using a time step of 2 fs.

For the tetrameric coiled-coil domain (residues C91 to D155), we additionally performed a simulation starting from the equilibrated structure (50 ns) at 310 K and heated up the system gradually to reach the temperature of 400 K and continued the simulation for 240 ns. The structure at the end of the simulation was used as the starting structure for the simulation with the temperature of 310 K, which was done for 680 ns.

For the HBD region together with the tetrameric coiled-coil as well as for the TMD inside the membrane, we additionally performed simulations under the electric field corresponding to the electric potential difference of 200 mV and 100 mV across the simulation cell, respectively. These simulations were performed for 1 µs and for systems with $Na^+$ and $Ca^{2+}$ ions.

#### Steered MD simulations

To check the stability of the tetrameric coiled-coil (residues C91 to D155), we applied harmonic force with a force constant of 1500 kJ $mol^{-1}$ $nm^{-2}$ to the center of mass of the opposite chains to increase the distance between them. The rate was chosen as 0.01 nm per ns. One structure was then selected in which the chains have been opened so that the alpha helices in the chains have not been distorted and then this structure was simulated for 1 µs.
To pull the $Ca^{2+}$ ion inside the HBD and subsequently into the tetrameric coiled-coil, a harmonic pulling force was applied to the ion with a force constant of 1500 kJ $mol^{-1}$ $nm^{-2}$ and the rate of 0.1 nm per ns opposite to the direction of the z-axis.

#### Metadynamics simulations

The metadynamics simulation was performed on the combination of the cryo-EM structure of prepore state, using residues C91 to D155 (one chain) and the corresponding coiled-coil region of the pore-state (three chains) predicted by AlphaFold2, employing version 2019.6 of GROMACS[22,23] and version 2.6.4 of Plumed[30]. The collective variable was chosen as the RMSD of the chain, taken from the cryo-EM structure, with respect to the corresponding chain, predicted from AlphaFold2. During the simulation several restrains were applied: 1) upper wall potential with the force constant of 1000 kJ $mol^{-1}$ $nm^{-2}$ to the RMSD with the maximum value of 2.1 Å. 2), lower wall potential with the force constant of 1000 kJ $mol^{-1}$ $nm^{-2}$ to the number of alpha helices. This number was constrained to be larger or equal to 235. 3) lower wall potential with a force constant of 1000 kJ $mol^{-1}$ $nm^{-2}$ to the distance between the beginning and end residue of the cryo-EM part of the structure with the lowest value of 3.8 nm. Gaussian hills with the height of 0.1 kJ/mol and width of 0.02 nm were added to the biasing potential every 1 ps. The final energy was constructed finally as the negative of the biasing potential.

#### Analysis MD data

The simulation data were analyzed using GROMACS tools as well as in-house codes in Python, incorporating MDAnalysis[31,32]. VMD[33] and ChimeraX[18,19] were used to visualize the structures and trajectories as well as to prepare the snapshots and the movies.
The pore volume as displayed in Figure 3 was created by moleonline[34] and to calculate the pore profiles of the proteins the HOLE program[35] was used. For all simulations the pore profile was calculated every $10^{th}$ ns and the average and the standard deviation of the pore were calculated and then represented with a solid line and a shaded area, respectively.
To calculate the ion permeation events, upper and lower thresholds corresponding to the z position of the Cα atoms of the S92 and G61 residues were considered, respectively, and then when an ion during the simulations had a continuous trajectory with the maximum z-position value higher than 5 Å above the upper threshold and the minimum z-position value lower than 5 Å below the lower threshold, the ions were considered as crossed and the trajectory of the ion was recorded.

To calculate the density of the ions along the xy-axis, the protein was considered at the center and the ion positions along the xy-axis were accordingly translated. The histograms were calculated for different slabs along the z-axis and averaged over the simulation time.
For the distance of the two adjacent helices, for all Cα atoms of one helix the minimum distance to a Cα atom from second helix is determined. The overall distance is given as the average value of these distances.
